# Turnover and replication analysis by isotope labeling (TRAIL) reveals the influence of tissue context on protein and organelle lifetimes

**DOI:** 10.1101/2022.04.24.488979

**Authors:** John Hasper, Kevin Welle, Jennifer Hryhorenko, Sina Ghaemmaghami, Abigail Buchwalter

## Abstract

The lifespans of proteins can range from minutes to years within mammalian tissues. Protein lifespan is relevant to organismal aging, as long-lived proteins can accrue damage over time. It is unclear how protein lifetime is shaped by tissue context, where both cell division and proteolytic degradation contribute to protein turnover. Here, we develop turnover and replication <u>a</u>nalysis by ^15^N isotope labeling (TRAIL) to quantify both protein and cell lifetimes with high precision and no toxicity over a 32-day labeling period across 4 mammalian tissues. We report that cell division promotes non-selective protein turnover in proliferative tissues, while physicochemical features such as hydrophobicity, charge, and intrinsic disorder exert a significant influence on protein turnover only in non-proliferative tissues. Protein lifetimes vary non-randomly across tissues after correcting for differences in cell division rate. Multiprotein complexes such as the ribosome have highly consistent lifetimes across tissues, while mitochondria, peroxisomes, and lipid droplets have variable lifetimes. These data indicate that cell turnover, sequence-encoded features, and other environmental factors modulate protein lifespan *in vivo*. In the future, TRAIL can be used to explore how environment, aging, and disease affect tissue homeostasis.

## Introduction

The cellular proteome undergoes constant cycles of synthesis, folding, and degradation. Proteostasis (protein homeostasis) is achieved by the balance of these processes. When these systems function properly, the health of the proteome is ensured by the efficient degradation of misfolded or damaged proteins and replacement with properly folded and functional copies. When proteostasis breaks down due to aging or disease, proteome disruptions including accumulation of oxidative damage, misfolding, and aggregation result^1,2^. Measurements of protein turnover have revealed that protein lifetimes range from minutes to years within mammalian tissues^3–7^. The functional consequences of age-linked proteostasis collapse are most evident for extremely long-lived proteins in post-mitotic tissues. For instance, crystallin proteins of the eye lens misfold and aggregate over decades, causing cataracts^8^, while the extremely long-lived nuclear pore complex becomes leaky and dysfunctional in the aging brain^9^. These striking examples raise several questions, including: what factors control protein lifetime in healthy tissues? What is the relationship between protein longevity and cellular longevity? Why do age-linked declines in long-lived protein function manifest only in some tissues?

Protein lifetime can be influenced by both sequence-encoded features and environmental factors^10^. For instance, proteins with long disordered segments are generally more short-lived than proteins that adopt a stable structure^11,12^. Post-translational modifications have varied effects on protein stability^13,14^, while higher buried surface area correlates with longer lifetime^15^. There are many exceptions that break these rules, however, and it is unclear to what extent physicochemical features predict protein lifetime *in vivo*. Additionally, the cellular, tissue and organismal environments of proteins can strongly influence their degradation rates. For instance, the same protein sequence can have dramatically different lifetimes when expressed in different cell types, tissues or organisms^5,16–19^.

One important environmental parameter that can strongly influence the observed turnover rate of a protein is the proliferative capacity of the tissue where it is expressed. Protein clearance (on a per cell basis) is influenced by the additive effects of its degradation kinetics as well as cellular dilution due to cell division^3,5^ (Fig. 1A). Thus, in general, protein clearance rates are expected to be faster within proliferative tissues in comparison to non-proliferative tissues. However, in typical dynamic metabolic labeling experiments employed for measurements of *in vivo* protein turnover, potential differences in tissue proliferation rates are typically unknown, making it impossible to deconvolute the effects of protein degradation and dilution. Thus, with currently available methodology, it is not possible to account for differences in tissue proliferation when comparing protein turnover rates across tissues.

**Figure 1.**
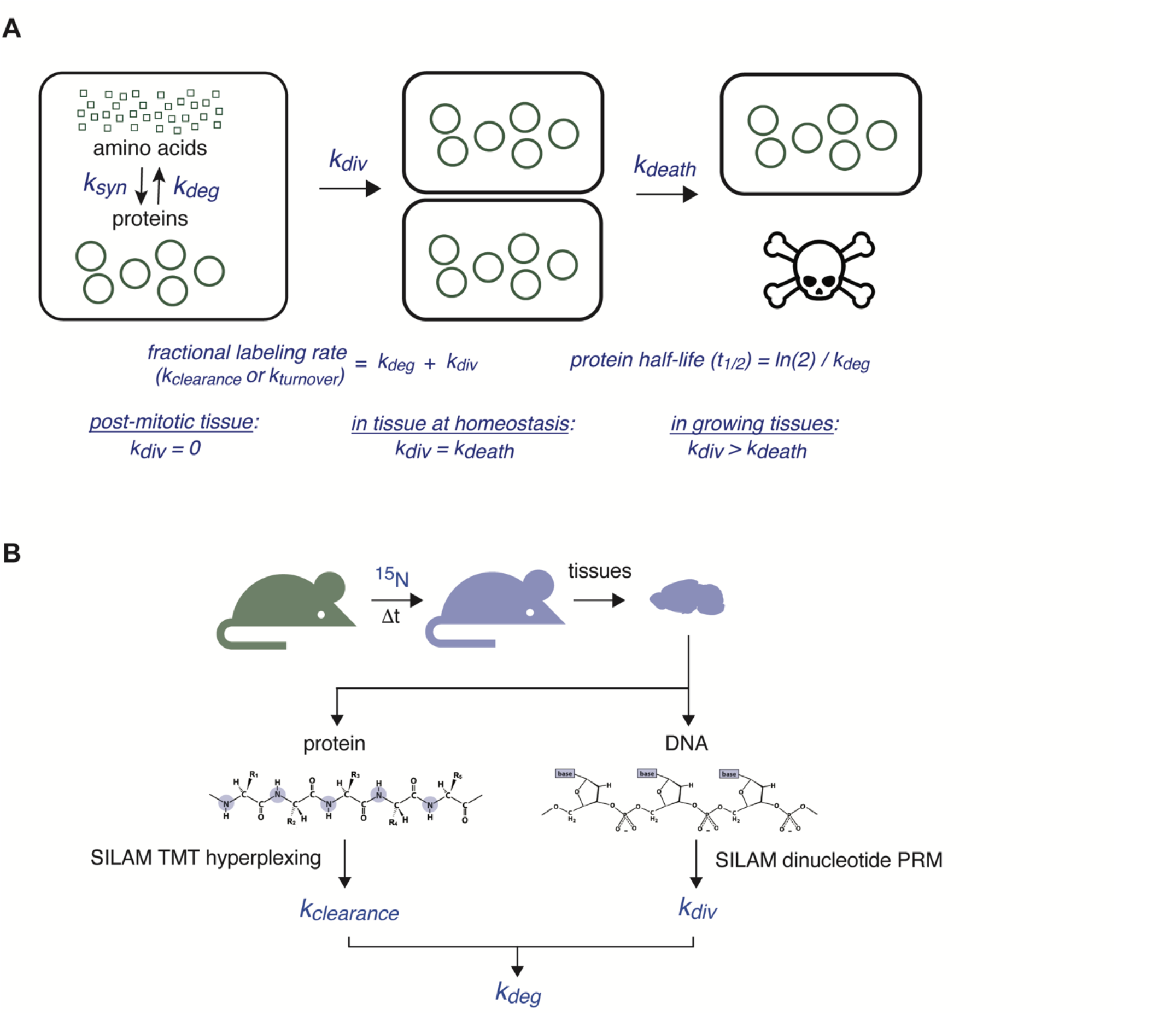
(A) The fractional rate by which a protein population is turned over within a cell (k_t_) can be determined by measuring the fractional rate of isotope incorporation in continuous labeling experiments. This rate us established by the additive effects of protein degradation (k_deg_) and cell division (k_div_)- In post-mitotic cells, k_div_ is negligible and does not contribute to protein turnover. In proliferating tissues, the rate of cell division is balanced by the rate of cell death (k_death_)-(B) Diagram of turnover and replication analysis by isotope labeling (TRAIL) approach for simultaneously quantifying k_t_ and k_div_ These two rates are measured by quantifying rates of ^15^N incorporation into proteins and DNA within the same experiment. Together, these two measurements can be used to accurately measyre k_deg_.

To accurately measure *in vivo* protein turnover rates within multiple tissues, we sought to develop a mass spectrometry-based method capable of simultaneously quantifying in vivo protein degradation and cell division rates within a single labeling experiment. While metabolic labeling with ^13^C or ^15^N isotopes has become the gold standard for the quantification of protein turnover rates^3,20^, this methodology has not been integrated with cell turnover measurements. Instead, cell turnover rates are frequently measured by partial labeling with nucleotide analogs (e.g. ^3^H-thymidine or BrdU), an approach that is often limited by label toxicity^21,22^. Alternatively, D_2_O labeling has been used to quantify cell turnover^23^ or to measure bulk rates of protein turnover and nucleic acid turnover ^24,25^. However, only low levels of D_2_O can be tolerated *in vivo*, and the small mass shifts that are achieved by partial labeling require specialized analysis methods for quantitation^26^. Here, we describe methods to measure both protein degradation and cell division within mammalian tissues using a single source of label: the stable isotope ^15^N (Fig. 1B). We name this suite of methods “turnover and replication <u>a</u>nalysis by isotope labeling”, or TRAIL. We apply TRAIL to proliferative and non-proliferative tissues and generate a rich dataset that reveals tissue-specific features of proteostasis. We find evidence for sequence-based selectivity in protein turnover in tissues that undergo slow cell proliferation, while protein turnover is much less selective in highly proliferative tissues. Further, protein and organelle lifetimes vary widely across healthy tissues even after correcting for cell proliferation rates. These observations illustrate the variable influence of “nature” (sequence-encoded features) *versus* “nurture” (environmental factors) on proteostasis *in vivo*. In the future, TRAIL can be used to explore how environment, aging, and disease affect tissue homeostasis.

## Results

### Increasing throughput of stable isotope labeling timecourses by tandem mass tag multiplexing

Proteome-wide quantification of protein stability can be achieved *in vivo* by feeding mice a food source containing ∼100% abundance of the stable, non-toxic isotope ^15^N, a method referred to as stable isotope labeling in mammals (SILAM)^3,20^. Labeled tissues are then analyzed by tandem mass spectrometry (LC-MS/MS) to quantify the incorporation of labeled amino acids into the proteome over time^3,20^. Broader application of SILAM has been limited by the investment of resources and time required to complete these types of analyses. One major bottleneck is mass spectrometer run time, which rapidly multiplies when each sample must be analyzed in a separate LC-MS/MS run. Further, protein “dropout” due to missing values limits the number of proteins whose turnover kinetics can be precisely determined. We have previously used tandem mass tagging (TMT) to “hyperplex” pools of isotope-labeled samples in a single LC-MS/MS run, which decreases cost while increasing speed and sensitivity^27^. Here, we adapt this approach to ^15^N-labeled samples from mouse tissue (TMT-SILAM, Fig. 2A). Hyperplexed analysis of ^15^N-labeled samples presents a unique challenge due to the high complexity of the labeled peptide spectra. The gradual labeling of the *in vivo* amino acid precursor pool by ^15^N results in broadened MS1 spectra whose average mass to charge ratios increase as a function of labeling time^3^, creating a challenging analysis problem that requires specialized data analysis workflows^28,29^. However, we previously demonstrated that hyperplexed sample analysis can be simplified by quantifying the relative decay of unlabeled peaks as a function of time rather than the fractional population of unlabeled and labeled peaks^27^. Here, by quantifying the fractional rate of loss of ^14^N peptides (as newly synthesized ^15^N labeled peptides accumulate), we were able to directly measure the turnover rate of pre-existing unlabeled proteins (Fig. 2A).

**Figure 2.**
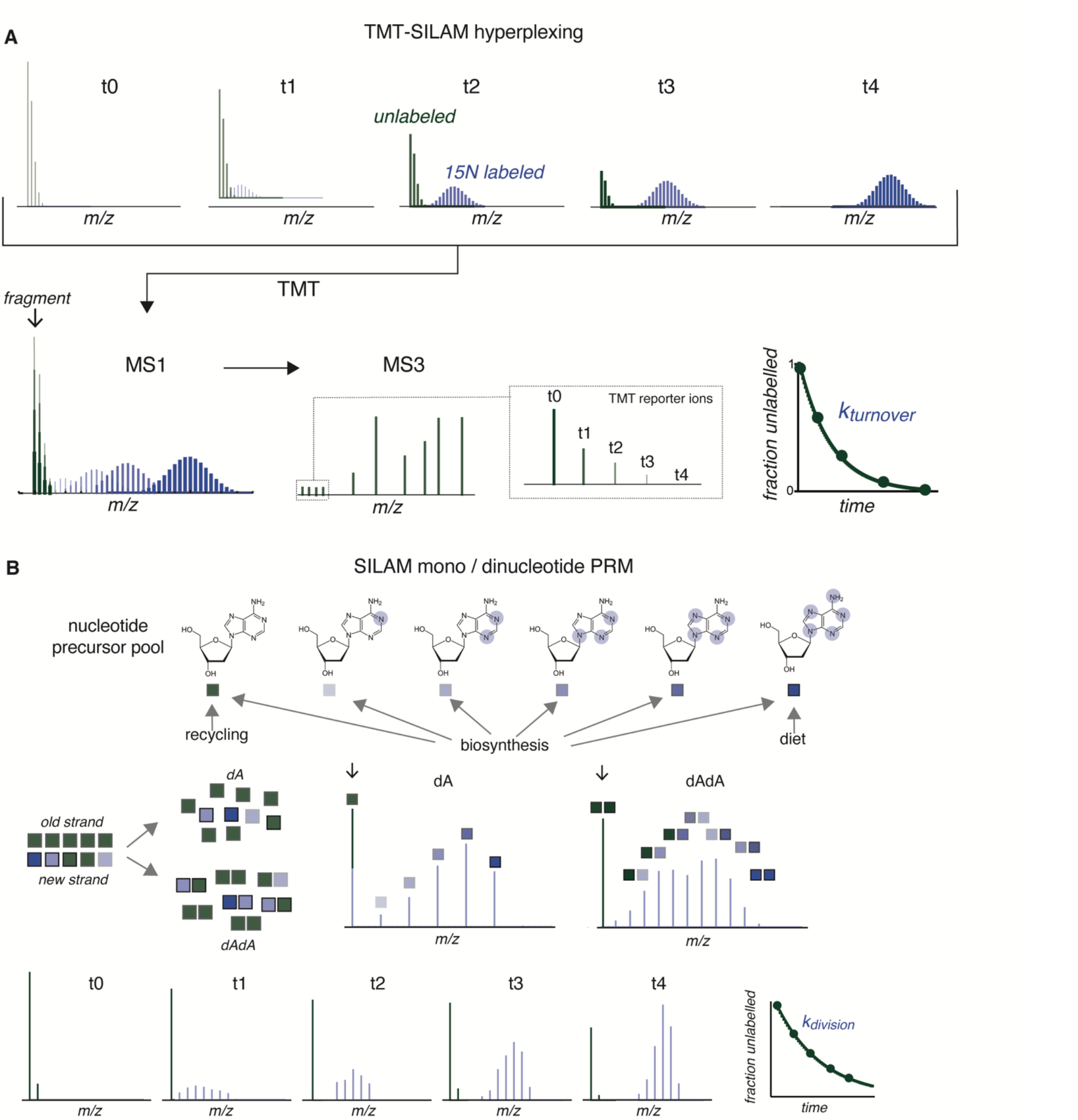
(A) Schematic of TMT-SILAM hyperplexed determination of *k*_*t*_ by tracking the rate of loss of unlabeled peptides over time. Samples from multiple timepoints are hyperplexed by conjugation with unique TMT tags and combined into a single sample before LC-MS/MS. A peptide detected in multiple time points will have a single m/z in MS1, but will fragment into reporter ions with distinct masses in MS2/MS3. Quantification of SILAM-unlabeled MS3 peaks reveals the kinetics of loss of the unlabeled peptides, which can be fit to determine *k*_*t*_. (B) Schematic of SILAM mono and dinucleotide PRM. The relative isotope abundance (RIA) of the nucleotide precursor pool reflects the rates of precursor uptake, biosynthesis, and recycling of pre-existing nucleotides. By quantifying the abundances of unlabeled, partially labeled, and fully labeled dinucleotides in genomic DNA, the extent of unlabeled nucleotide recycling can be inferred and can be used to determine corrected cell division rates (*k*_*div*_).

We performed a 32-day TMT-SILAM time course on young adult (9-week-old) mice. Animal weights remained stable through the labeling period (Supplementary Figure 1), indicating that protein levels are at a steady-state and fractional labeling rates can be equated with protein turnover rates^30^ (see Methods). We focused our analyses on selected tissues that are thought to be either highly proliferative or largely post-mitotic^31^: the large intestine (a proliferative tissue); the liver (a quiescent tissue that can proliferate in response to injury); and the heart and white adipose tissue, which are mostly post-mitotic. We analyzed the labeling kinetics of thousands of proteins per tissue (Supplementary Figure 2) and filtered these data at several levels to compile high quality datasets. Experimental replicates were first filtered based on coverage: only proteins that were detected with a minimum of 3 peptide spectral matches (PSMs) in all channels were retained for further analysis. Second, aggregated replicate data were used to determine the rate constant for protein turnover (*k*_*t*_) by least squares fitting to a first order kinetic model (Note that *k*_*t*_ values refer to protein turnover rate constants that have not been corrected for the dilution effects of cell division as described below). Only *k*_*t*_ values that were measured by fitting data arising from at least 2 replicates with a high goodness of fit (t-statistic > 3, see Methods) were considered in downstream analyses.

### Features of protein turnover across tissues

Altogether, we defined high-confidence *k*_*t*_ values (Fig. 3A) and corresponding predicted half-life (*t*_*1/2*_) values (Fig. 3B) for thousands of proteins per tissue: 2719 in the large intestine, 2099 in liver, 1610 in white adipose tissue, and 1635 in the heart (Supplementary Fig. 2; Supplementary Table 1). Protein abundance and protein lifetime were generally not correlated with each other (Fig. 4A; Supplementary Fig. 4). The distributions of *k*_*t*_ values were unique for each tissue; proteins were more short-lived in the intestine (median t_1/2_ 1.7 days) and liver (median t_1/2_ 2.4 days), but more long-lived in the fat (median t_1/2_ 6.1 days) and heart (median t_1/2_ 5.7 days). Differences in protein stability have been previously reported between mammalian tissues, such as the brain, liver, and muscle^3,5,19^. We compared our datasets to previous analyses of proteome turnover in the liver^3^ and heart^32^ and found high concordance in both cases (Supplementary Figure 5).

**Figure 3.**
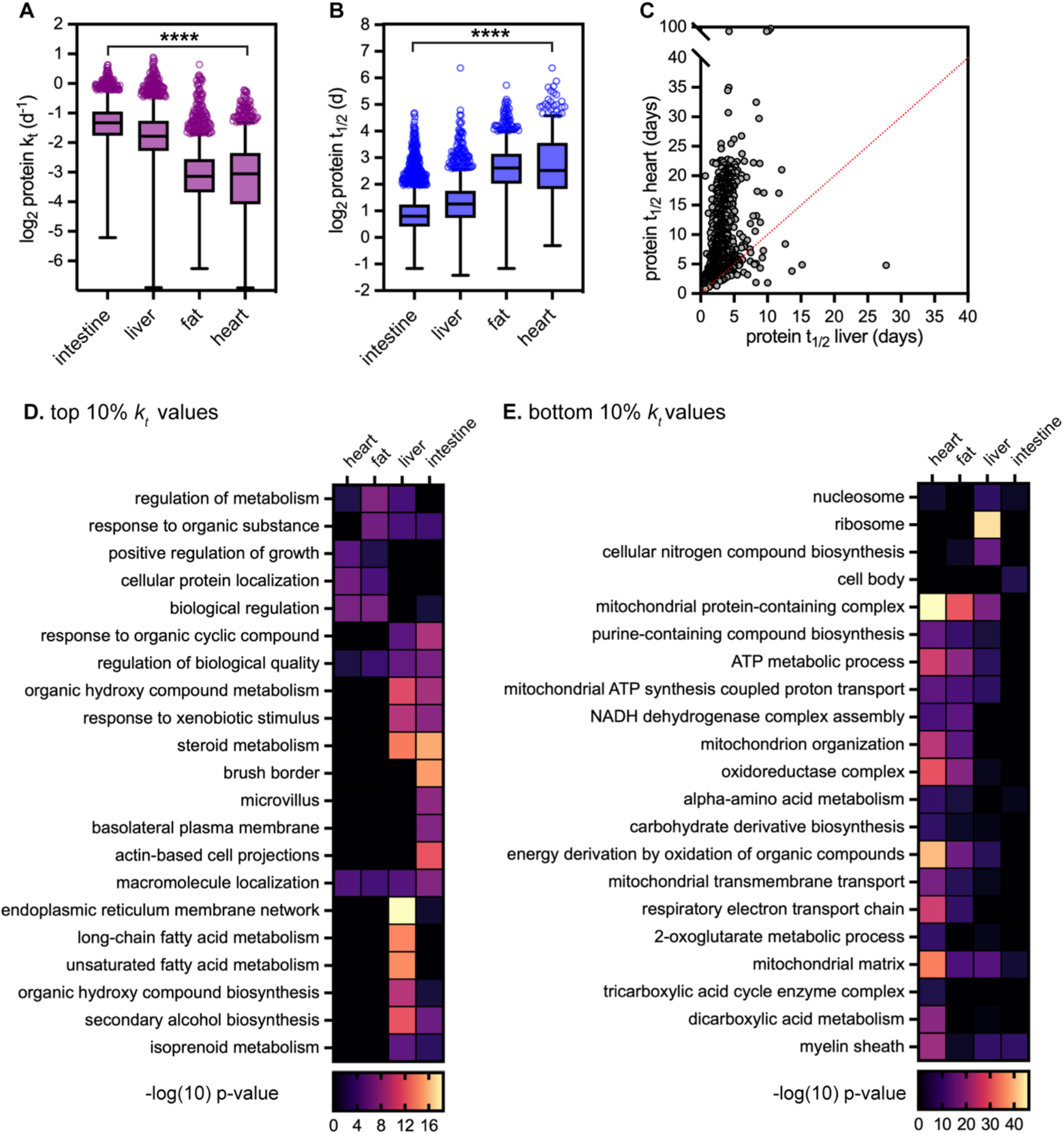
Proteome turnover measurements for four reference tissues. (A) Turnover rates (*k*_*t*_) and (B) predicted half-lives (t_1/2_) determined by SILAM-TMT of proteins extracted from intestine (n = 2719; median *t*_1/2_ 1.7 days), liver (n = 2099, median *t*_1/2_ 2.4 days), fat (n = 1610, median *t*_1/2_ 6.1 days), and heart (n = 1635, median *t*_1/2_ 5.7 days). Box (Tukey) plot center line indicates median; box limits indicate 25th to 75th percentiles; whiskers indicate 1.5x interquartile range; points indicate outlier values. **** indicates that all medians are significantly different (p < 0.0001, Kruskal-Wallis test). See also Supplementary Table 1 for full dataset. (C) Predicted half-lives (*t*_1/2_) for 1102 intracellular proteins in the heart versus the liver. (D-E) Heatmaps of GO term enrichment in the top 10% (least stable, D) and the bottom 10% (most stable, E) of *k*_*t*_ values for intracellular proteins detected in each tissue.

**Figure 4.**
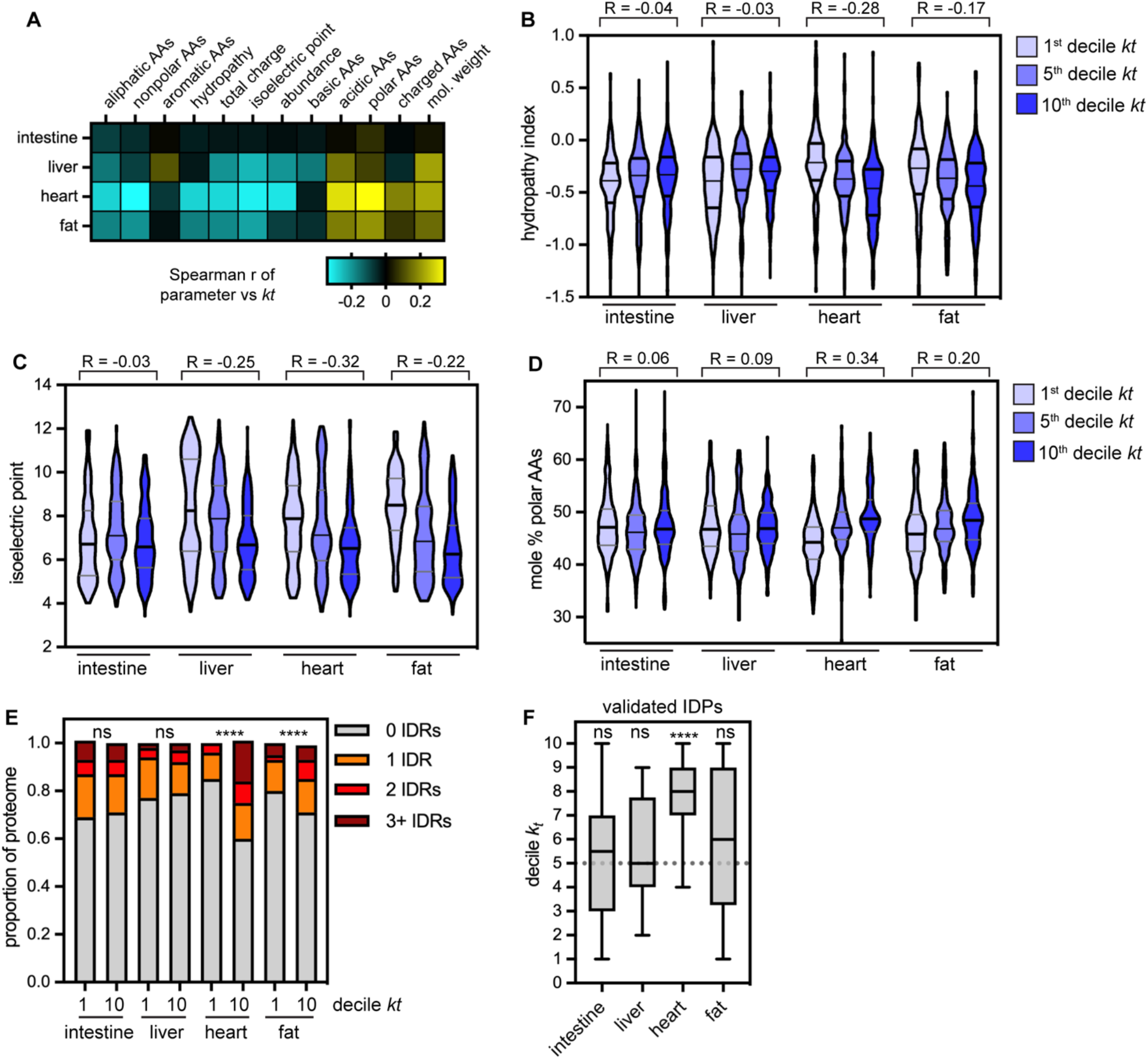
Analysis of correlations between protein sequence features and protein turnover rate across tissues. (A) Heatmap of Spearman correlation coefficient between *kt* and protein sequence features (see Methods). (B) GRAVY hydropathy index of proteins in the 1st, 5th, and 10th *kt* deciles in each tissue; hydropathy is significantly anticorrelated with *kt* in heart and fat proteomes, but not in intestine or liver proteomes. (C) Isoelectric point of proteins in the 1st, 5th, and 10th *kt* deciles in each tissue; pl is significantly anti-correlated with *kt* in the liver, heart, and fat proteomes, but not in the intestine proteome. (D) Abundance of polar and charged amino acids (mole% of D, E, H, K, N, Q, R, S, T) in the 1st, 5th, and 10th *kt* deciles in each tissue; polar / charged residue abundance is significantly positively correlated with *kt* in the heart and fat proteomes, but not in the intestine or liver proteomes. (E) Incidence of proteins containing long (> 40 AA) intrinsically disordered regions (IDRs) in the 1st and 10th *kt* deciles in neach tissue. IDR-containing proteins are significantly over-represented in the most short-lived proteins in the heart and fat proteomes, but not in the intestine or liver proteomes. **** indicates p < 0.0001; significance determined by χ2 test. (F) Turnover rates (*kt* decile) of 16 experimentally validated IDPs are significantly faster than the proteome median in the heart, but not in other tissues. IDP annotations from the DisProt database.

We explored and excluded several potential explanations for tissue-specific differences in protein stability. (1) These differences persist among the subset of proteins whose lifetimes were quantified across all four tissues (902 proteins total; Supplementary Fig. 6A-B; Supplementary Table 2), indicating that differences in protein lifetime are not due simply to differences in proteome composition. Further, a large proportion of protein lifetime differences are statistically significant in pairwise comparisons between tissues, while 282 proteins (31%) have significantly different *k*_*t*_ values across all four tissues (Supplementary Fig. 6C; Supplementary Table 2). (2) These differences also persist if secreted proteins are excluded from analyses (Supplementary Fig. 6D-F); secreted proteins are a unique class of proteins whose turnover is difficult to accurately profile *in vivo*, as their sites of synthesis, function, and degradation can be quite disparate. Even after controlling for these factors, differences in protein lifetimes across tissues are readily apparent. For example, many intracellular proteins have days-long lifetimes in the liver but weeks-long lifetimes in the heart (Fig. 3C), with 85% of these proteins having significantly different *k*_*t*_ values between these two tissues (Supplementary Fig 6C).

We looked more closely at intracellular proteins at the extremes of stability by identifying gene ontology (GO) terms that were over-represented in either the top decile (Fig. 3D) or bottom decile (Fig. 3E) of *k*_*t*_ values in each tissue. The identity of the most short-lived and most long-lived proteins varies widely. For instance, proteins involved in intestinal nutrient absorption are enriched among the most short-lived proteins of the large intestine, while proteins involved in alcohol and fatty acid metabolism are found among the most short-lived proteins of the liver (Fig. 3D). At the other extreme, components of chromatin are enriched in the most long-lived proteins of the liver and intestine, while proteins involved in various mitochondrial functions are enriched in the most long-lived proteins of the heart and adipose tissue (Fig. 3E). Interestingly, while components of chromatin and mitochondria have been found to be long-lived in other protein turnover studies^3,5,33,34^, our data suggest that their relative stability varies from tissue to tissue.

We next explored how physicochemical properties such as amino acid composition, hydrophobicity, charge, and intrinsic disorder correlate with protein turnover (Fig. 4; Supplementary Table 3). While these relationships have been explored within protein turnover datasets acquired in yeast^35^ and in mammalian cell culture^11,12,36^, they have not to our knowledge been evaluated across mammalian tissues. Overall, we did not find a single protein feature that correlated significantly with protein turnover rate across all tissues. Rather, we found features that showed significant relationships to protein turnover in a subset of tissues (Fig. 4A). For instance, hydrophobicity decreases as turnover rate increases in the heart and fat proteomes, but not in the liver or intestine proteomes (Fig. 4A-B). In these same tissues, polar amino acids are more abundant in short-lived proteins than in long-lived proteins (Fig. 4A, 4D). Protein isoelectric point is strongly anti-correlated with *k*_*t*_, such that long-lived proteins are more basic (pI > 7) while short-lived proteins are more acidic (pI < 7) in the heart, liver, and fat (Fig. 4A, 4C). Consistently, acidic amino acids are over-represented in short-lived proteins in these tissues (Fig. 4A). Finally, we evaluated relationships between protein disorder and protein turnover by quantifying the frequency of intrinsically disordered regions (IDRs) in the most stable and least stable proteins (Fig. 4E). IDRs of at least 40 amino acids in length are correlated with significantly accelerated protein turnover across eukaryotes^11,12^. While IDRs are over-represented in unstable heart and fat proteins, there is no relationship between disorder and protein lifetime in the intestine or liver (Fig. 4E). In an orthogonal approach, we evaluated the turnover rates of an experimentally validated list of disordered proteins from the DisProt database^37^ and found that this validated group of disordered proteins turned over significantly faster than the proteome median within the heart, but not in any other tissue (Fig. 4F; Supplementary Fig. 6G-H). Instead, many of these disordered proteins are rapidly degraded in one tissue but relatively stable in another. Taken together, our analyses reveal greater sequence-based selectivity of turnover of the heart and fat proteome than of the liver and intestine proteome.

We speculate that the interplay between sequence features and environmental factors influences protein lifetime *in vivo*. As described above, within a living tissue, the rates of protein turnover are influenced both by proteolytic degradation of proteins and by dilution of proteins during cell division (Fig. 1A). Variations in the extent of cell proliferation could, at least in part, underlie the observed differences in protein turnover rates across tissues (Fig. 3). Differences in cell proliferation may also influence the observed differences in distributions of protein turnover rates within a given tissue (Fig. 4) as cell division non-selectively accelerates apparent protein turnover rates among all proteins within a given tissue. To accurately define the relationship between protein lifetime and cellular proliferation, a method that can quantify both of these parameters in parallel is needed.

### Turnover and replication analysis by isotope labeling (TRAIL) to profile cell and protein turnover

We sought to develop a method to quantify cell proliferation rates in parallel with protein turnover measurements. It is well appreciated that ^15^N-labeled nutrients supplied via SILAM chow can efficiently label proteins in mice. However, ^15^N can also be robustly incorporated into genomic DNA *via* nitrogen-containing nucleobases^38^. We therefore reasoned that tracking the rate of ^15^N incorporation into the genome via DNA replication would yield cell division rates (*k*_*div*_), which when conducted in conjunction with analyses of protein labeling could be used to determine corrected protein degradation rates (*k*_*deg*_)^30^ (Fig. 1B, Fig. 2). We refer to this method as turnover and replication <u>a</u>nalysis by isotope labeling (TRAIL).

To develop this approach, we first needed to address the technical barrier imposed by the prevalence of in vivo nucleotide recycling. The fractional labeling of genomic DNA during an isotope labeling experiment is influenced both by the rate of replication and by the relative isotope abundance (RIA) of the precursor nucleotide pool (Fig. 2B). The latter is strongly influenced by precursor uptake from the diet, nucleotide biosynthesis, and nucleotide recycling *in vivo*^23^. Incomplete labeling of the precursor pool due to low precursor uptake, slow *de novo* biosynthesis, or extensive recycling of pre-existing nucleic acids would decrease the extent of ^15^N incorporation into replicating genomic DNA^32^. Further, the relative contributions of each of these factors may vary across tissues. Thus, it is important to define the RIA of the precursor nucleotide pool in each tissue in order to accurately determine the rate of replication by measuring the fractional labeling of genomic DNA. We reasoned that we could deconvolute the RIA of the precursor nucleotide pool by analyzing the combinatorics of labeling in contiguous stretches of *dinucleotides* obtained from the same strand of genomic DNA (Fig. 2B, Supplementary Fig. 8; see Methods). A strand of DNA that has been synthesized in the presence of label will contain labeled nucleotides at a frequency that is contingent on the RIA of the precursor pool. Analysis of the isotopologue distribution within dinucleotides enables the calculation of the prevalence of recycled unlabeled nucleotides within the precursor pool. Thus, we can determine what fraction of the observed fully unlabeled dinucleotide population was derived from pre-existing unlabeled DNA strands, and what fraction was derived from newly synthesized strands that incorporated unlabeled recycled nucleotides. Through this deconvolution, we can measure the relative ratio of old and newly synthesized DNA and determine the rate of cell proliferation. We digested genomic DNA from SILAM-labeled mouse tissue to short oligonucleotides using the enzyme benzonase^39^, then quantified ^15^N/^14^N isotope abundance ratios in dinucleotides by mass spectrometry. We found that the ^15^N RIA of the precursor pool was very high in all tissues, and that diet-derived ^15^N-labeled nucleic acids were preferentially incorporated into newly synthesized genomic DNA (Supplementary Fig. 8C). This is in line with the fact that nucleotide salvage pathways are repressed in S phase while *de novo* nucleotide synthesis is upregulated, so that cells primarily rely on the latter source of nucleotides for DNA replication^40^. This outcome is also consistent with observations from other modes of DNA labeling^41^.

Our finding that diet-derived and de novo synthesized nucleic acids are preferred for DNA replication implies that we can make an accurate measurement of DNA replication rates by tracking ^15^N incorporation into either mononucleosides or dinucleotides isolated from genomic DNA. We tested this by isolating free dA, dC, dT, and dG mononucleosides from genomic DNA by digestion with a cocktail of benzonase, phosphodiesterase, and alkaline phosphatase^42^ and quantifying ^15^N/^14^N isotope ratios by mass spectrometry (see Methods). Decay curves for all four mononucleosides were in close alignment both with each other (Supplementary Fig. 9A) and with dinucleotide curves (Supplementary Fig. 8D), indicating that TRAIL is highly precise and reproducible.

To test how accurately TRAIL reports cell division rates, we devised a benchmarking experiment as follows. We isolated fibroblasts from the ear of a mouse that had undergone ^15^N labeling for a total of 256 days. Because mouse fibroblasts renew within weeks, the genomic DNA from these cells was highly labeled with ^15^N. We then subcultured these fibroblasts *ex vivo* and collected genomic DNA at 3 timepoints over the course of several days. In parallel, we quantified cell numbers. We then compared the doubling times determined by TRAIL *versus* direct measurement of population doublings. These data were highly consistent (Supplementary Figure 10), indicating that TRAIL accurately tracks cell division rate.

With these important controls established, we then applied TRAIL to the large intestine. We determined a doubling time of ∼3 days for this proliferative epithelial tissue, in close agreement with previous analyses by orthogonal methods^31,43^ (Fig. 5A-B). We then applied this method to the liver, fat, and heart and found that cells of these tissues have very long average doubling times indicating low proliferative capacity (Fig. 5A-B; Supplementary Table 4). These data are qualitatively consistent with previous reports of slow proliferation in these tissues^44–46^(Fig. 5A-B). Notably, however, analysis of isotope incorporation into both nucleosides and dinucleotides provides far greater precision and accuracy than methods that infer cell doubling time from pulse-labeling with a single nucleotide analog (such as ^3^H-thymidine or BrdU) or immunostaining with a cell-cycle-linked antigen (such as Ki-67).

**Figure 5.**
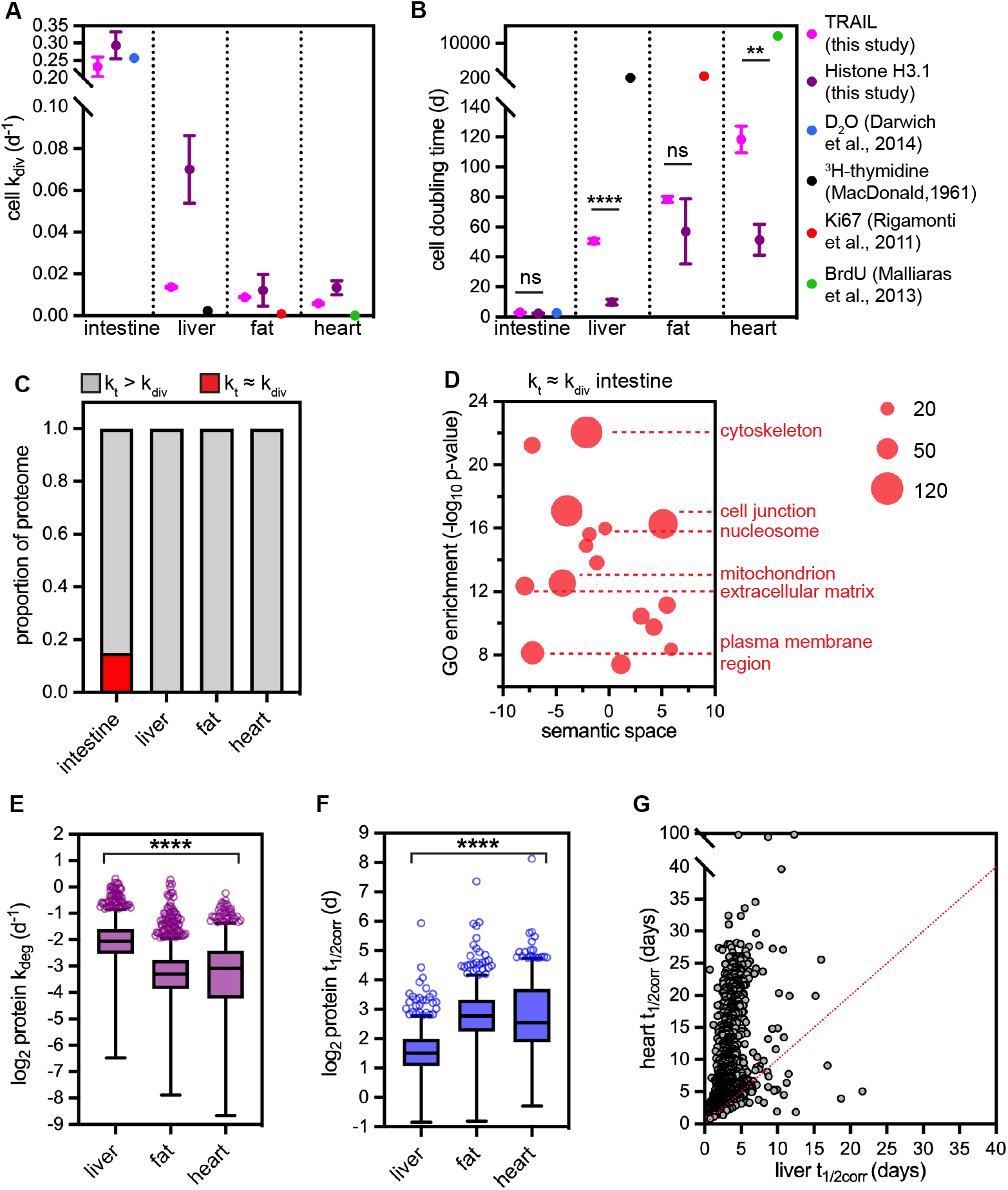
(A-B) Comparison of cell division rate (*k*_*div*_, (A)) and cell doubling time (B) in intestine, liver, fat, and heart, determined by TRAIL (our study); by Histone H3.1 protein turnover (our study); and previously published reports by a range of methods, including 3H-thymidine labeling, BrdU pulse-labeling, D_2_ O water labeling, and Ki67 staining. Error bars indicate SEM. Data reproduced from meta-analysis by Sender & Milo Nature Medicine 2021. * indicates p < 0.05 for *k*_*div*_ determined by TRAIL versus H3.1 turnover in the liver (unpaired t-test). See also Supplementary Table 3 for full dataset. (C) Only the proliferative intestine has a significant number of proteins whose *k*_*t*_ is equivalent to *k*_*div*_, suggesting that these proteins are diluted by cell division. (D) A subset of Gene Ontology (GO) terms (cellular component) over-represented in proteins that are diluted out by cell division in the intestine are shown in a bubble plot. Redundant GO terms were removed and non-redundant GO terms were organized based on semantic similarity by REViGO. Bubble size corresponds to number of proteins associated with GO term. (E) Cell-cycle-corrected degradation rates (*k*_*deg*_) and (F) half-lives (*t*_*1/2corr*_) for 967 proteins detected in liver (median *t*_*1/2*_ 2.9 days), fat (median *t*_*1/2*_ 6.8 days), and heart (median *t*_*1/2*_ 5.9 days). Box (Tukey) plot center line indicates median; box limits indicate 25th to 75th percentiles; whiskers indicate 1.5x interquartile range; points indicate outlier values. **** indicates that all medians are significantly different (p < 0.0001, Kruskal-Wallis test). (G) Predicted cell-cycle-corrected half-lives (*t*_*1/2corr*_) for 1102 intracellular proteins in the heart versus the liver. See also Supplementary Table 4 for full dataset.

The turnover of Histone H3.1 has been used as a proxy for cell division in protein turnover studies^5,16^ because Histone H3.1 is incorporated into nucleosomes as a heterodimer with Histone H4 solely after DNA replication^47^, leading to the expectation that this protein’s levels would decrease by dilution over successive cell divisions. However, we noted that the lifetime of Histone H3.1 was often shorter than the cell doubling time determined by TRAIL. This distinction was most apparent in the slowly proliferative liver, heart, and fat (Fig. 5A-B). We speculate that this difference reflects DNA replication-independent processes that regulate the lifetime of H3.1. For instance, H3.1/H4 dimers can be evicted from DNA during transcription and are replaced with heterodimers of Histone H3.3 and Histone H4^48^. Separately, histones can also be found in cytosolic pools in complex with chaperones, where they may be more rapidly turned over^49^. We speculate that each of these factors contributes to the turnover of this histone isoform in postmitotic tissues over long timescales. TRAIL thus provides a superior measurement of cell proliferation rates over long timescales and suggests contextual variability in the rate of turnover of the replication-dependent Histone H3.1 across tissues.

### Protein degradation rates vary across tissues after cell cycle correction

A powerful feature of TRAIL is the ability to co-capture cell and protein turnover from the same tissues. Accordingly, cellular *k*_*div*_ rates can be subtracted from protein *k*_*t*_ values to extrapolate corrected protein degradation rates (*k*_*deg*_)^30^ (Fig. 1A). The extreme case where a *k*_*deg*_ value approaches zero indicates that dilution by cell division is the major contributor to protein turnover; that is, protein *k*_*t*_ and cellular *k*_*div*_ are approximately equal. When we compared these values across tissues, we found that *k*_*div*_ was roughly equivalent to *k*_*t*_ for approximately 15% of the intestine proteome (Fig. 5C; Supplementary Fig. 9B). These proteins are components of diverse cellular structures including the cytoskeleton, the extracellular matrix, cell-cell junctions, mitochondria, and chromatin (Fig. 5D). Notably, many of these proteins are also long-lived in postmitotic tissues but turn over at a slow rate that exceeds the rate of cell division. Histone H3.1 is one such protein (Fig. 5A-B). This observation implies that long-lived proteins are non-selectively diluted by cell division in highly proliferative tissues but are slowly and selectively turned over in less proliferative tissues.

We next examined *k*_*deg*_ values in the slowly proliferative liver, heart, and fat. If variable cell proliferation rates underlie the variability in protein *k*_*t*_ values across these tissues, *k*_*deg*_ values should be largely invariant after correcting for *k*_*div*_. We did not observe this outcome. Instead, the range of *k*_*deg*_ values remained distinct from tissue to tissue even after correcting for cell proliferation rates, and persisted when we restricted our analysis only to proteins that were detected in all tissues (967 proteins; Fig. 5E-F; Supplementary Table 5-6). Overall, the liver proteome (median *t*_*1/2corr*_ 2.9 days) turns over significantly more rapidly than the fat proteome (median *t*_*1/2corr*_ 6.8 days; 90% of *k*_*deg*_ values are significantly different) or heart proteome (median *t*_*1/2corr*_ 5.8 days; 84% of *k*_*deg*_ values are significantly different) after cell cycle correction (Fig. 5G). These data indicate that protein lifetime is broadly influenced by other environmental factors beyond cell proliferation rate. Consistent with our findings, protein lifetimes have also been found to differ significantly between non-dividing cell types in culture^16^, as well as in the same cell type (fibroblasts) isolated from different mammals^17^. One potential explanation for these differences could be variation in the composition and activity of protein folding chaperones, the ubiquitin-proteasome system, and/or the autophagy machinery across tissues^50–52^.

### Peroxisomes, lipid droplets, and mitochondria have highly variable lifetimes across tissues

To evaluate the extent of variability in *k*_*deg*_ across tissues, we determined the normalized cross-tissue dispersion (*D)* of *k*_*deg*_ for the 967 proteins shared across the liver, fat, and heart datasets (see Methods; Supplementary Fig. 11; Supplementary Table 7). We then used this metric to dissect variability in protein lifetime across tissues, focusing on constituents of cellular organelles (Fig. 6), multiprotein complexes (Fig. 7) and pathways (Supplementary Fig. 12).

**Figure 6.**
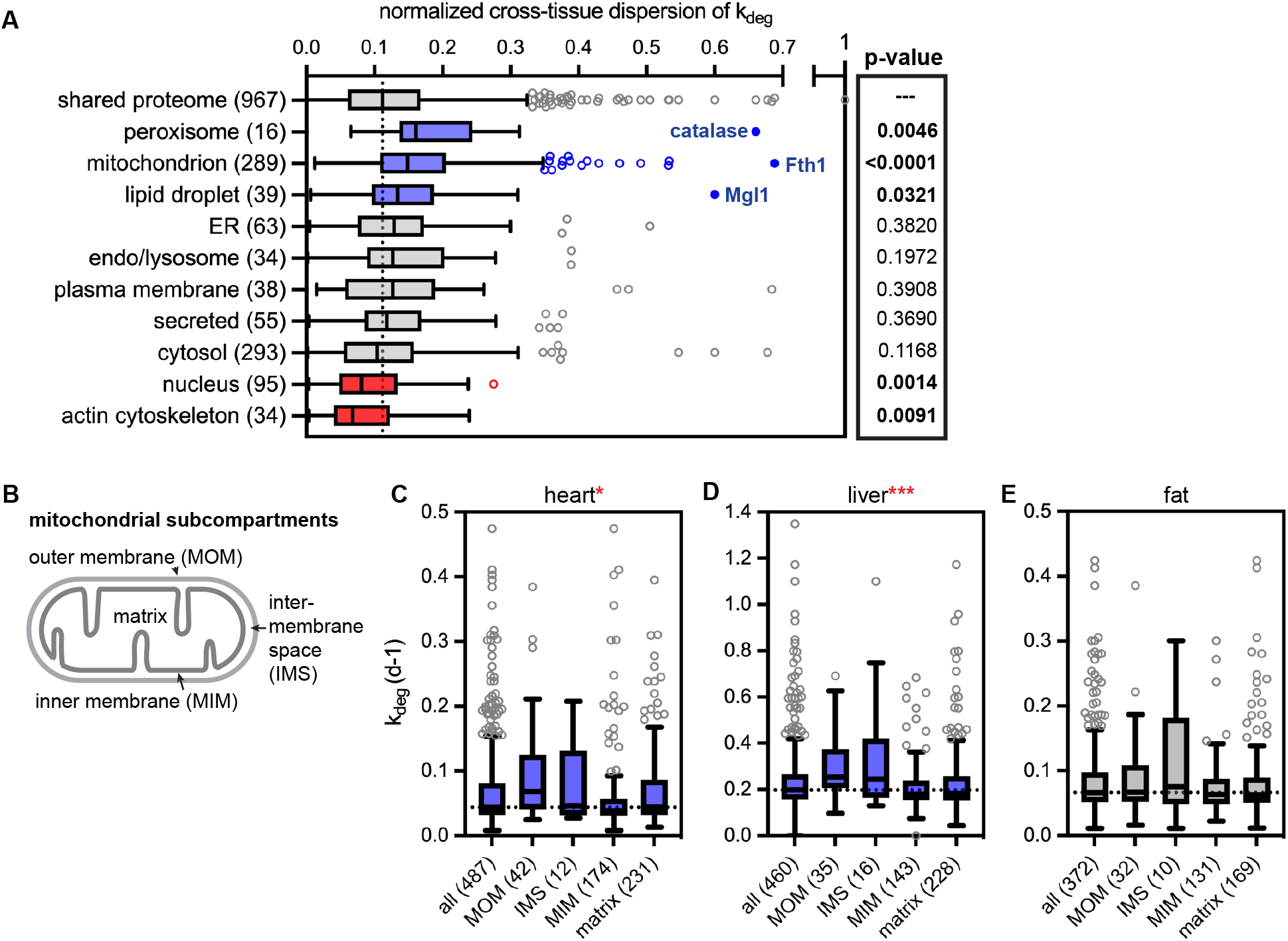
(A) Cross-tissue dispersion analysis of *k*_*deg*_ values across heart, liver, and fat by cellular organelle. Blue indicates a subset has a significantly elevated median dispersion of *k*_*deg*_ values across tissues; red indicates a subset has a significantly decreased dispersion of *k*_*deg*_ values across tissues. Significance determined by Mann-Whitney test. The peroxisomal enzyme catalase, the mitochondrial resident protein Fth1, and the lipid droplet enzyme MGL1 are highlighted as residents of each organelle with the most extremely variable lifetimes across tissues. (B-D) Analysis of *k*_*deg*_ rates across mitochondrial sub-compartments. MOM: mitochondrial outer membrane; IMS: intermembrane space; MIM: mitochondrial inner membrane. Numbers indicate total proteins detected in each subcompartment. Proteins of the MOM and IMS turn over significantly faster than proteins of the MIM and matrix in the heart (C) and liver (D). In contrast, all mitochondrial subcompartments turn over at similar rates in the white adipose tissue (E). Significance determined by Kruskal-Wallis test.See also Supplementary Table 5 for full dataset. All box (Tukey) plots: center line indicates median; box limits indicate 25th to 75th percentiles; whiskers indicate 1.5x interquartile range; points indicate outlier values.

**Figure 7.**
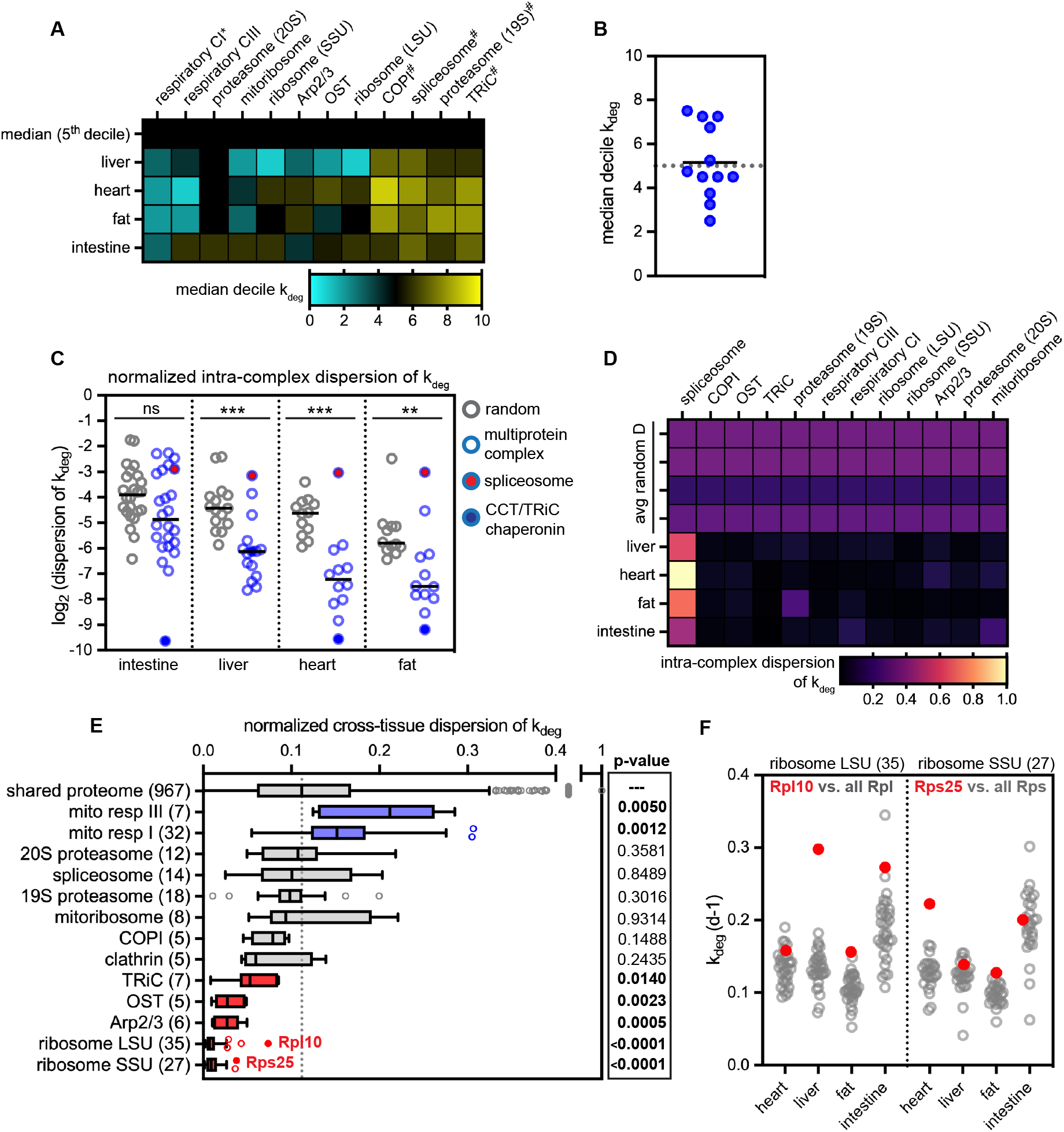
(A) The median decile of *k*_*deg*_ for each multiprotein complex with at least 5 subunits detected. *indicates complexes that turn over significantly more slowly, and #indicates complexes that turn over significantly faster than the proteome median in all tissues. (B) Median decile *k*_*deg*_ for multiprotein complexes (black bar) does not deviate from the proteome median (5th decile). (C) Intra-complex dispersion was computed for complexes with at least 5 subunits detected (blue) and a random dispersion value was calculated by computing dispersion for an equivalent number of randomly chosen proteins (gray).Black bar, median. In liver, heart, and fat, complexes exhibit a significantly lower intra-complex dispersion than would be expected by chance (** p<0.01; *** p < 0.001, Mann-Whitney test). The spliceosome (red) is an outlier with high intra-complex dispersion; the TriC/chaperonin complex (solid blue) is an outlier with low intra-complex dispersion. (D) heatmap of intra-complex dispersion across tissues. (E) Analysis of normalized cross-tissue dispersion of *k*_*deg*_ by multiprotein complex. P-values indicate significance of deviation from proteome mean (Mann-Whitney test). While the small and large subunits of the ribosome have extremely low cross-tissue dispersion, outlier subunits Rps25 and Rpl10 have more variability across tissues (solid red). Box (Tukey) plot center line indicates median; boxes indicate 25th to 75th percentiles; whiskers indicate 1.5x interquartile range; points indicate outlier values. (F) *k*_*deg*_ values of ribosome subunits. See also Supplementary Table 5 for full dataset.

Among proteins that are residents of specific subcellular organelles, components of the actin cytoskeleton and residents of the nucleus have much less variable lifetimes than the proteome as a whole (Fig. 6A-B). In contrast, constituents of peroxisomes, lipid droplets, and mitochondria have significantly more variable lifetimes across tissues than the proteome as a whole (Fig. 6A; Supplementary Fig. 12), implying that the degradative flux of these organelles varies from tissue to tissue. These organelles can be degraded by specialized variants of autophagy termed pexophagy^53^, lipophagy^54^, and mitophagy^55^, respectively.

Peroxisomes play major roles in lipid catabolism, and their biogenesis is induced by signaling through peroxisome proliferator agonist receptors (PPARs) and other mechanisms. Upon removal of biogenesis-promoting signals, excess peroxisomes are degraded by pexophagy^56^. This process was first documented in the liver; consistently, we observe rapid turnover of peroxisomal proteins in this tissue. Our data indicate that peroxisomes are also degraded rapidly in the intestine but are degraded more slowly in the heart and adipose tissue (Supplementary Fig. 13).

Lipid droplets (LDs) are the major intracellular sites of lipid storage. In response to nutrient deprivation, LDs mobilize lipids either by lipolysis to generate fatty acids or by lipophagy, which involves delivery of both the protein and lipid components of LDs to the lysosome^57^. Lipophagic flux is high in the liver^54^, and we observe rapid degradation of LD proteins in this organ (Supplementary Fig. 13). In contrast,

LD proteins are longer-lived in the white adipose tissue and in the heart. This is somewhat unexpected, as adipose tissue rapidly mobilizes free fatty acids when nutrients are low^58^, while heart tissue depends on fat oxidation for energy^59^. This outcome indicates that lipophagic flux is lower in these tissues and suggests that fatty acids are instead mobilized from LDs by lipolysis while sparing LD resident proteins from turnover.

Mitochondria are long-lived organelles in many tissues, including the brain^3,6^, heart^32^, and skeletal muscle^33,34^. Consistent with these recent studies, we find that mitochondria are long-lived in the heart (median mitochondrial protein *t*_*1/2*_ of 18.1 days vs. 5.8 days for total proteome). Mitochondria are also long-lived in the white adipose tissue (median mitochondrial protein *t*_*1/2*_ of 10.5 days vs. 6.8 days for total proteome) (Fig. 6C,E). Surprisingly, however, mitochondria turn over more rapidly in the liver (median mitochondrial protein *t*_*1/2*_ of 3.5 days vs. 2.9 days for total proteome) (Fig. 6D). This finding suggests major differences in mitochondrial regulation and function in this organ but is consistent with a previous report of high mitophagy flux in the liver using an *in vivo* reporter system^60^.

We achieved high coverage of the mitochondrial proteome, making it possible to inspect the turnover of mitochondrial subcompartments across tissues (Fig. 6B-E). Proteins of the mitochondrial outer membrane (MOM) and the intermembrane space (IMS) generally turn over more rapidly than proteins of internal compartments such as the mitochondrial inner membrane (MIM) and the matrix. This disparity is most apparent in the heart and liver (Fig. 6C,D). A previous analysis of protein lifetimes in the brain similarly reported more rapid turnover of MOM proteins^6^. Notably, the MOM and IMS are accessible to the cytosol while the MIM and matrix are sequestered. These data suggest that selective degradation of MOM/IMS proteins occurs at a significant rate in many tissues, including the heart, liver, and brain^6^. This could be achieved by extraction and delivery to the proteasome, piecemeal autophagy, and/or sequestration into mitochondrial-derived vesicles^61^. In contrast, all mitochondrial subcompartments exhibit coherent turnover in white adipose tissue (Fig. 6E). This could indicate that mitochondria are degraded more frequently by organellar autophagy (mitophagy) than by selective degradation of mitochondrial components in this tissue. Interestingly, mitophagy plays major roles in the differentiation and maintenance of white adipocytes, which characteristically have lower numbers of mitochondria^62^. Altogether, our data indicate that both the overall flux of mitochondrial turnover and the mechanisms used to achieve turnover vary across tissues.

### Multiprotein complex subunits have coherent lifetimes within and across tissues

It has been suggested that participation in stable multiprotein complexes might protect proteins from degradation and extend protein half-life^15,63^. We evaluated the degradation kinetics of 12 multiprotein complexes for which at least 5 subunits were detected in all four tissues and found that in general, multiprotein complex subunits do not exhibit significantly lower degradation rates than the proteome median (Fig. 7A-B). This indicates that participation in a stable multiprotein complex is not sufficient to dramatically extend protein lifetime. It is important to note, however, that this steady-state measurement cannot determine whether complex subunits are selectively degraded if they fail to assemble correctly after synthesis^63^.

Some multiprotein complexes have been reported to exhibit coherent subunit turnover, perhaps reflecting their stable association from biogenesis to degradation^7^. To determine whether complex subunits exhibit more similar turnover rates than would be expected by random chance, we calculated the intra-tissue *k*_*deg*_ dispersion *(d)* for multiprotein complexes for which at least 5 subunits were detected (12-24 complexes per tissue). For comparison, we calculated *d* for an equivalent number of randomly chosen proteins (Fig. 7C,D). Comparing these values indicated that most multiprotein complexes turn over coherently in the liver, heart, and fat, while this difference was not significant in the intestine. Consistent with a previous report^7^, we also find that the CCT/TriC chaperonin complex is an outlier whose subunits have extremely consistent *k*_*deg*_ values (low *d*, Fig. 7C, solid blue). Other multiprotein complexes with coherent turnover include the ribosome, proteasome, oligosaccharyltransferase (OST) complex, and the mitochondrial respiratory chain complexes (Fig. 7D). In striking contrast, the spliceosome is an outlier whose components have widely varying *k*_*deg*_ values (high *d*; Fig. 6C, solid red; Fig. 7D). We speculate that this may reflect the cyclical, activity-coupled nature of spliceosomal assembly^64^, and may indicate that individual spliceosome subcomplexes undergo quality control / degradation at various rates.

We next asked whether multiprotein complex lifetimes are consistent or variable across tissues by calculating the cross-tissue *k*_*deg*_ dispersion *(D)* of individual subunits across the liver, heart, and fat. Components of mitochondrial respiratory chain complexes were the only complex subunits that had significantly higher *D* than the proteome median (Fig. 7E), which is likely due to the dramatic differences in mitochondrial lifetime across tissues (Fig. 6). Apart from these outliers, other multiprotein complex subunits had average or significantly lower than average values of *D*. We noted that the small and large subunits of the ribosome had extremely low *D* values (Fig. 7E), and that the stability of the small and large subunits tracked very closely with each other (Fig. 7F). The ribosome also has a very consistent half-life across fibroblasts derived from a range of mammalian species^17^, indicating that ribosome turnover is very tightly controlled. However, Rpl10 and Rps25 had much more variable *k*_*deg*_ values than other ribosomal proteins (Fig. 7E,F). Interestingly, Rpl10 association is a key late regulatory step in large subunit biogenesis^65^, and Rps25 is incorporated only in a subset of ribosomes that are endowed with unique translational specificity^66^. Rpl10 turns over faster than other large subunit components in the liver and intestine, while Rps25 turns over faster than most small subunit components in the heart (Fig. 7F). These data suggest nodes of ribosome biogenesis control that vary across tissues.

## Discussion

Here we report the development of TRAIL, a multiplexed ^15^N isotope labeling workflow that enables simultaneous measurements of protein lifetime and cellular lifetime from the same tissue. To our knowledge, this is the first study to advance a method for deriving cell turnover rates from ^15^N labeling. In contrast to other frequently used approaches for quantifying cell turnover, ^15^N has no detectable toxicity, even through multiple generations of continuous labeling in mice^4,20^, opening the possibility of extending TRAIL to accurately define the turnover rates of slowly proliferating cell types.

By sensitively measuring cell turnover and protein turnover in parallel, TRAIL adds a critical layer of context to analysis of proteostasis. We have unambiguously determined that protein lifetimes vary widely across tissues, and that sequence features as well as cell turnover and additional environmental factors shape protein lifetime. Our data suggest that long-lived proteins experience a very different life cycle in post-mitotic *versus* proliferative tissues. We found that dilution by cell division contributes non-selectively to the turnover of about 15% of the proteome in the proliferative intestine, such that these proteins are renewed roughly every 3 days as the epithelium renews (Figure 5). In contrast, protein lifetimes spread over a broader dynamic range in post-mitotic tissues, where protein turnover is both sequence-selective (Figure 4) and coordinated across multiprotein complex subunits (Figure 7). We speculate that only in this context would long-lived proteins and complexes meaningfully “age” - meaning that they accumulate oxidative damage, misfold, and lose their function, which would in turn lead to age-linked tissue dysfunction. We have also uncovered evidence that the rate of organelle degradation, perhaps by autophagy of peroxisomes, lipid droplets, and mitochondria, varies widely across tissues (Figure 6). Why do some proteins and organelles turn over at such variable rates? It is possible that protein damage occurs more rapidly in some tissues than in others, perhaps linked to the variable rate of production of reactive oxygen species and other damaging agents during normal cellular metabolism. A second, non-exclusive possibility is that the activity and/or selectivity of protein folding and/or degradation machineries varies across cell types and tissues^51^. Intriguingly, *in vivo* reporters of the proteasome and of autophagy do suggest variable flux across tissues^50,52^.

We anticipate that TRAIL can be applied to explore the consequences of aging and disease on tissue homeostasis. However, there are some limitations of our approach to consider. Our continuous labeling approach must assume maintenance of homeostatic balance over the time frame of the experiment-an assumption that is more likely to be valid over shorter timescales and in healthy tissues, but may not prove to be true over longer timescales or in diseased tissues. Our approach also does not address the contribution of different cell types to bulk measurements of cell turnover or protein turnover. We surmise that our data most accurately reflect the turnover of proteins that are broadly expressed in most cell types of the tissues analyzed. Overall, we were able to profile the 1500-3000 most abundant proteins per tissue, which tend to be broadly expressed and not cell-type-specific. Nevertheless, tissue atlases generated by single cell sequencing can provide a useful framework for thinking about the likely cell types of origin for the proteins profiled in this study^67^. Future studies may involve computational deconvolution or sorting of individual abundant cell types from tissues of interest in order to generate cell type-resolved maps of cell and proteome lifetime.

## Methods

### Metabolic labeling of mice and tissue isolation

We designed a 6-timepoint, 32-day SILAM labeling timecourse (0, 2, 4, 8, 16, and 32 days of labeling) with a total of 3 animals of both sexes per labeled timepoint, and 2 animals for the day 0 (unlabeled) timepoint. Timecourses were performed in male and female wild type C57Bl/6 mice at approximately 9 weeks of age. ^14^N and ^15^N mouse chow was obtained from Silantes. Animals were first habituated to the chow formulation by feeding ^14^N (normisotopic) food for 1 week and monitoring animal weight. Animals maintained normal weight through the duration of the timecourse. Animals were then transitioned to ^15^N chow throughout the labeling period (roughly 3 grams / animal / day). Animals were then sacrificed by CO_2_ inhalation followed by cervical dislocation, followed by tissue dissection and flash freezing by submersion in liquid nitrogen. These animal experiments were performed in compliance with relevant ethical regulations and with approval by the Institutional Animal Care and Use Committee at UCSF (IACUC protocol number AN178187, PI: A.B.).

### Protein extraction and sample preparation for LC-MS/MS

#### Protein extraction

Approximately 30 milligrams of frozen tissue was excised on dry ice with a clean razorblade and placed in a fresh tube. 100 uL of protein extraction buffer (PEB: 5% SDS, 100 mM TEAB, protease and phosphatase inhibitors, pH ∼7) was added to the tube. The tissue was rapidly minced with clean dissection scissors on ice for 30-60 seconds until no large pieces remained. PEB was added to bring the final volume to 600 uL, then the sample was transferred to a Dounce homogenizer. The sample was homogenized for ∼40 strokes with the tight pestle, then was transferred to a clean microcentrifuge tube. The sample was then probe sonicated at 4C (10% amplitude, 10 seconds, 2 cycles) before being centrifuged (21,000 x g, 11 minutes, 4C). The supernatant was transferred to a clean tube, and aliquots were separated for proteomics and protein quantification by microBSA assay (Pierce).

#### Trypsinization

Samples were diluted to 1 mg/mL in 5% SDS, 100 mM TEAB, and 25 μg of protein from each sample was reduced with dithiothreitol to 2 mM, followed by incubation at 55°C for 60 minutes. Iodoacetamide was added to 10 mM and incubated in the dark at room temperature for 30 minutes to alkylate the proteins. Phosphoric acid was added to 1.2%, followed by six volumes of 90% methanol, 100 mM TEAB. The resulting solution was added to S-Trap micros (Protifi), and centrifuged at 4,000 x g for 1 minute. The S-Traps containing trapped protein were washed twice by centrifuging through 90% methanol, 100 mM TEAB. 1 μg of trypsin was brought up in 20 μL of 100 mM TEAB and added to the S-Trap, followed by an additional 20 μL of TEAB to ensure the sample did not dry out. The cap to the S-Trap was loosely screwed on but not tightened to ensure the solution was not pushed out of the S-Trap during digestion. Samples were placed in a humidity chamber at 37°C overnight. The next morning, the S-Trap was centrifuged at 4,000 x g for 1 minute to collect the digested peptides. Sequential additions of 0.1% TFA in acetonitrile and 0.1% TFA in 50% acetonitrile were added to the S-trap, centrifuged, and pooled. Samples were frozen and dried down in a Speed Vac (Labconco) prior to TMTpro labeling.

#### TMT labeling

Samples were reconstituted in TEAB to 1 mg/mL, then labeled with TMTpro 16plex reagents (Thermo Fisher) following the manufacturers protocol. Briefly, TMTpro tags were removed from the -20°C freezer and allowed to come to room temperature, after which acetonitrile was added. Individual TMT tags were added to respective samples, and the reaction was allowed to occur at room temperature for 1 hour. 5% hydroxylamine was added to quench the reaction, after which the samples for each experiment were combined into a single tube. Since we performed quantitation on the unlabeled peptides, 0 day samples were added to four of the unused channels, increasing the signal for the unlabeled peptides. TMTpro tagged samples were frozen, dried down in the Speed Vac, and then desalted using homemade C18 spin columns to remove excess tag prior to high pH fractionation.

#### High pH Fractionation

Homemade C18 spin columns were activated with two 50 μL washes of acetonitrile via centrifugation, followed by equilibration with two 50 μL washes of 0.1% TFA. Desalted, TMTpro tagged peptides were brought up in 50 μL of 0.1% TFA and added to the spin column. After centrifugation, the column was washed once with water, then once with 10 mM ammonium hydroxide. Fractions were eluted off the column with centrifugation by stepwise addition of 10 mM ammonium hydroxide with the following concentrations of acetonitrile: 2%, 3.5%, 5%, 6.5%, 8%, 9.5%, 11%, 12.5%, 14%, 15.5%, 17%, 18.5%, 20%, 21.5%, 27%, and 50%. The sixteen fractions were concatenated down to 8 by combining fractions 1 and 9, 2 and 10, 3 and 11, etc. Fractionated samples were frozen, dried down in the Speed Vac, and brought up in 0.1% TFA prior to mass spectrometry analysis.

### LC-MS/MS Analysis

*Data collection:* Peptides from each fraction were injected onto a homemade 30 cm C18 column with 1.8 um beads (Sepax), with an Easy nLC-1200 HPLC (Thermo Fisher), connected to a Fusion Lumos Tribrid mass spectrometer (Thermo Fisher). Solvent A was 0.1% formic acid in water, while solvent B was 0.1% formic acid in 80% acetonitrile. Ions were introduced to the mass spectrometer using a Nanospray Flex source operating at 2 kV. The gradient began at 3% B and held for 2 minutes, increased to 10% B over 7 minutes, increased to 38% B over 94 minutes, then ramped up to 90% B in 5 minutes and was held for 3 minutes, before returning to starting conditions in 2 minutes and re-equilibrating for 7 minutes, for a total run time of 120 minutes. The Fusion Lumos was operated in data-dependent mode, employing the MultiNotch Synchronized Precursor Selection MS3 method to increase quantitative accuracy^68^. The cycle time was set to 3 seconds. Monoisotopic Precursor Selection (MIPS) was set to Peptide. The full scan was done over a range of 400-1500 m/z, with a resolution of 120,000 at m/z of 200, an AGC target of 4e5, and a maximum injection time of 50 ms. Peptides with a charge state between 2-5 were picked for fragmentation. Precursor ions were fragmented by collision-induced dissociation (CID) using a collision energy of 35% and an isolation width of 1.0 m/z. MS2 scans were collected in the ion trap with an AGC target of 1e4 and a maximum injection time of 35 ms. MS3 scans were performed by fragmenting the 10 most intense fragment ions between 400-2000 m/z, excluding ions that were 40 m/z less and 10 m/z greater than the precursor peptide, using higher energy collisional dissociation (HCD). MS3 ions were detected in the Orbitrap with a resolution of 50,000 at m/z 200 over a scan range of 100-300 m/z. The isolation width was set to 2 Da, the collision energy was 60%, the AGC was set to 1e5, and the maximum injection time was set to 100 ms. Dynamic exclusion was set to 45 seconds.

#### Data analysis

Raw data was searched using the SEQUEST search engine within the Proteome Discoverer software platform, version 2.4 (Thermo Fisher), using the Uniprot mouse database (downloaded January 2020). Trypsin was selected as the enzyme allowing up to 2 missed cleavages, with an MS1 mass tolerance of 10 ppm, and an MS2 mass tolerance of 0.6 Da. Carbamidomethyl on cysteine, and TMTpro on lysine and peptide N-terminus were set as a fixed modifications, while oxidation of methionine was set as a variable modification. Percolator was used as the FDR calculator, filtering out peptides which had a q-value greater than 0.01. Reporter ions were quantified using the Reporter Ions Quantifier node, with an integration tolerance of 20 ppm, and the integration method being set to “most confident centroid”. Protein abundances were calculated by summing the signal to noise of the reporter ions from each identified peptide, while excluding any peptides with an isolation interference of greater than 30%, or SPS matches less than 65%.

#### Kinetic model

The kinetic model applied in this study has been previously described^27^. Briefly, we are assuming that protein synthesis is a zero order process, occurs at a constant fractional rate, and that that the total protein concentration of each cell does not change during the experimental time-course. The dilution of the protein pool due to cell division can be modeled as a first order exponential process. Thus, the fractional turnover of unlabeled proteins during the labeling time course can be regarded as a first order kinetic process that can be modelled based on the following exponential equation:

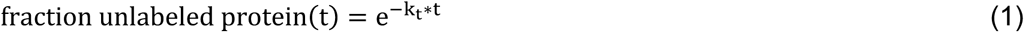

And:

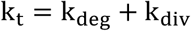

Where:

k_t_ is the clearance rate (observed rate of fractional labeling), k_deg_ is the rate of protein degradation and k_div_ is the rate of cell division.

The determination of k_t_ values were conducted as previously described^27^ using the decay of the TMT reporter signals of unlabeled proteins. Protein-level TMT reporter abundances for unlabeled proteins for each replicate experiment were first normalized by dividing by the intensity of the t0 reporter and then the replicate experiments were aggregated in a single kinetic curve. In fitting the exponential decay curves of the unlabeled protein signals, a constant fractional baseline at infinite time was incorporated in the fitting equation. The equation used for fitting the curves was therefore: 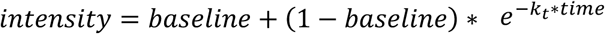. The goodness of fit for least squares fits were assessed by determining the R^2^, P-value and t-statistic of the fits (see Supp Table 1). For subsequent analyses, only protein k_t_ measurements that were obtained from all three replicate experiments, incorporated data from 4 or more peptide spectral matches (PSMs), and had t-statistic values greater than three were considered.

### Nucleic acid extraction and sample preparation for LC-MS/MS

#### Genomic DNA extraction

Approximately 30 milligrams of frozen tissue was excised on dry ice with a clean razorblade and placed in a fresh tube. 100 uL of TRIzol reagent (Invitrogen) was added and the tissue was rapidly minced with clean dissection scissors on ice for 30-60 seconds until no large pieces remained. An additional 400 uL of TRIzol was added, and the sample was then transferred to a Dounce homogenizer. The tissue was subjected to ∼40 strokes with the tight pestle until smooth, then transferred back to the original tube. The sample was incubated for at least 5 minutes before the addition of 100 uL chloroform followed by mixing and a further 3 minutes of incubation. The sample was then centrifuged (12,000 x g, 15 minutes, 4C) and the upper RNA-containing aqueous layer was discarded. 150 uL of absolute ethanol was added to the remaining sample, then inverted several times to mix. After 3 minutes of incubation at room temperature, the sample was centrifuged (2,000 x g, 5 minutes, 4C). The protein-containing supernatant was removed, then the DNA-containing pellet was resuspended in 500 uL of absolute ethanol and incubated for 30 minutes. The sample was then centrifuged (2,000 x g, 5 minutes, 4C), and the supernatant discarded. Sequential washes were then repeated with 95%, 85%, and 75% ethanol, after which the pellet was air-dried for 5-10 minutes. The pellet was then resuspended in 200 uL nuclease-free water (Ambion) at 56C, then incubated at 56C with shaking for 30 minutes to resuspend the pure DNA. The sample was centrifuged (12,000 x g, 10 minutes, 4C), then the supernatant containing pure DNA was moved to a clean tube. DNA concentration was determined with a NanoDrop spectrophotometer.

#### Digestion of genomic DNA to short oligonucleotides

3-5 micrograms of pure genomic DNA was diluted to a 50 uL volume in nuclease-free water, then combined with 50 uL of 2x Dinucleotide Buffer (DB: 5 mU/uL benzonase, 40 mU/uL shrimp alkaline phosphatase, 20 mM Tris pH 7.9, 100 mM NaCl, 20 mM MgCl2). Samples were incubated overnight at 37C. Spin-X UF Concentrators (Corning) were rinsed with 200 uL buffer (20 mM Tris pH 7.9, 100 mM NaCl, 20 mM MgCl2), then samples were applied and centrifuged through (12,000 x g, 5 min, RT). The eluate was collected for analysis.

#### Digestion of genomic DNA to mononucleosides

We extracted mononucleosides from genomic DNA similarly to a previously described method^42^ with some modifications. 1-3 micrograms of pure genomic DNA was diluted to a 50 uL volume in nuclease-free water, then combined with 50 uL of 2x Mononucleoside Buffer (MB: 5 mU/uL benzonase, 40 mU/uL shrimp alkaline phosphatase, 60 uU/uL phosphodiesterase I, 20 mM Tris pH 7.9, 100 mM NaCl, and 20 mM MgCl2). Samples were incubated overnight at 37C. Spin-X UF Concentrators (Corning) were rinsed with 200 uL buffer (20 mM Tris pH 7.9, 100 mM NaCl, 20 mM MgCl2), then samples were applied and centrifuged through (12,000 x g, 5 min, RT). The eluate was collected for analysis.

### Mononucleoside and Dinucleoside LC-MS/MS

Mononucleotide analyses were carried out by adapting a previously described method^69^ using a Dionex Ultimate 3000 UHPLC coupled to a Q Exactive Plus mass spectrometer (Thermo Scientific). After purification, analytes were separated on a Hypersil Gold 2.1 × 150 mm column, protected by a 2.1 × 10 mm Hypersil Gold guard column (Thermo Scientific). The mobile phases were A: 0.1% formic acid in water, and B: 0.1% formic acid in acetonitrile. The flow rate was set to 400 μL/min, and the column oven was set to 36°C. 10 μL of each sample was injected, and the analytes were eluted using the following gradient: 0 min- 0% B, 6 min- 0% B, 8.5 min- 80% B, 9.5 min- 80% B, 10 min- 0% B, 13 min- 0% B. The Q Exactive Plus was operated in positive mode with a heated electrospray ionization (HESI) source. The spray voltage was set to 3.5 kV, the sheath gas flow rate was set to 40, and the auxiliary gas flow rate set to 7, while the capillary temperature was set to 320°C. A parallel reaction monitoring (PRM) method was used to quantify the unlabeled nucleotide, along with all of its N15 isotopes in a single scan. This was accomplished by using wide (8 m/z) isolation widths when selecting the nucleotides for fragmentation. By employing this method, we were able to quantify the level of labeling by looking at the intensity of each N15 labeled base in the MS2 scan. Fragment ions were detected in the Orbitrap with a resolution of 70,000 at m/z 200. Using a high resolution MS2 scan allowed us to resolve N15 and C13 isotopes. Peak areas from the fragment ions were extracted with a 10 ppm mass tolerance using the LC Quan node of the XCalibur software (Thermo Scientific).

Dinucleotide analyses were carried out using the same instrumentation, column, mobile phases, column temperature, and flow rate employed by the mononucleotide experiments. The gradient was changed to optimize dinucleotide separation as follows: 0 min- 5% B, 0.5 min- 5% B, 2.5 min- 90% B, 3.25 min- 90% B, 3.5 min- 5% B, 5.5 min- 5% B. The Q Exactive Plus was operated using the same tune settings as the mononucleotide experiment. However, instead of a PRM method, a full scan method from 500-650 m/z was developed to quantify the dinucleotides dCdC, TT, dAdA, and dGdG, along with their corresponding N15 isotopes. Precursor ions were detected in the Orbitrap with a resolution of 140,000 at m/z 200, using the high resolution MS1 scan to try to separate N15 and C13 isotopes as much as possible. Peak areas from the fragment ions were extracted with a 10 ppm mass tolerance using the LC Quan node of the XCalibur software (Thermo Scientific).

#### Measurement of kdiv

To accurately measure rates of cell division (*k*_*div*_) while factoring in the effects of incomplete labeling and nucleotide recycling, we considered the time-dependent labeling patterns of mononucleotides and dinucleotides derived from genomic DNA. Upon initiation of ^15^N labeling, newly synthesized DNA strands can incorporate nucleotides from a precursor pool with potentially complex mixture of partially labeled species (Supplementary Fig. 8A). For example, a newly incorporated deoxyadenosine (dA) can be derived from fully ^15^N-labeled nucleotides derived from the dietary source, partially labeled species (containing one to four ^15^N atoms) derived by biosynthesis from incompletely labeled ^15^N precursors, and completely unlabeled nucleotides derived from recycling. As an example, a typical labeling pattern for dA from one of our intestine samples is shown in Supplementary Fig. 8B showing the shift in the isotopologue distribution over time. After correcting for the natural isotopic distribution, the peaks with heavier non-monoisotopic masses (+1, +2, +3, etc.) can be assumed to have been derived from newly synthesized strands. However, the monoisotopic peak (0) can potentially have been derived from both the original unlabeled strand, as well as newly synthesized strands that had incoroporated recycled unlabeled nucleotides. Therefore, it may not be possible to accurately determine the ratio of new to old strands (and hence *k*_*div*_) from the mononucleotide data alone. The labeling pattern of dinucleotides (dAdA) resolves this ambiguity. The isotopologue distribution of labeled (non-monoisotopic) peaks in the dinucleotides spectra are dependent on the composition of the nucleotide precursor pool. The red envelope depicted in the dAdA spectra is the pattern that would be expected if the precursor pool was composed solely of the labeled (non-monoisotopic) species observed in the corresponding mononucleotide spectra (i.e. new strands did not contain any recycled unlabeled nucleotides and the monoisotopic peaks observed in the mononucleotide spectra were derived solely from old strands). If a significant fraction of the monoisotopic peaks observed in the mononucleotide spectra represents recycled nucleotides within new strands, then the isotopologue distribution of labeled nucleotides would shift accordingly (Supplementary Fig. 8C). Through regression analyses, we determined that within all tissues and timepoints analyzed in this study, the isotopologue distributions of the dinucleotide data could be best modeled based on the assumption that newly synthesized strands had very low levels of fully unlabeled nucleotides. Hence, the fractional population of labeled non-monoisotopic peaks within dinucleotide and mononucleotide data were consistent with each other (Supplementary Fig. 8D) and could be used to determine the fractional population of new strands. For each tissue, fractional labeling of mononucleotide and dinucleotides for all 4 bases were combined and the aggregated dataset was fit to a single exponential equation to determine first order rate constant for division (*k*_*div*_). These data appear in Supplementary Table 4.

### Analysis of Proteomic Data

#### Quality filtering and analysis of k_*t*_ *values*

Proteomic data was acquired in the form of 16-plex TMT replicates containing 2 full 6-timepoint timecourses (one from wild type animals and one from progeroid animals). For each genotype within each TMT replicate, proteins were filtered to retain only those detected with at least 3 peptide spectral matches (PSMs) in all timepoints. Proteins that met these criteria were then filtered per genotype within each TMT replicate based on goodness of fit using the t-statistic. The t-statistic is equal to the turnover rate (*k*_*t*_) dividied by the standard error of that value. This metric determines to what extent measurement error influences *k*_*t*_. We applied a minimum t-statistic cutoff of 3, meaning that the magnitude of the turnover rate *k*_*t*_ is at least 3 times the magnitude of the standard error. Between 50% to 63% of detected proteins passed these coverage and goodness-of-fit criteria (Supplementary Figure 2). Along with the sample size, the t-statistic can be used to determine a p-value that indicates the probability that the turnover rate reported has a meaningful non-zero value. The *k*_*t*_, standard error, t-statistic, and p-value for each protein are reported in Supplementary Table 1. The *k*_*t*_, standard error, and sample size were used to perform per-protein statistical tests across tissues, to identify proteins with significantly different turnover kinetics between tissues. These data are reported in Supplementary Table 1.

Filtered *k*_*t*_ values for each tissue were separated into deciles. Proteins in the top decile (fastest *k*_*t*_) and bottom decile (slowest *k*_*t*_) were subjected to gene ontology analysis to identify biological processes (GO:BP) and cellular components (GO:CC) that were over-represented, using the Gprofiler tool^70^. Redundant GO terms were filtered using ReVIGO^71^, then subjected to hierarchical clustering and presented in heatmap format with cell values corresponding to the significance of enrichment for each term.

#### Analysis of protein sequence feature correlations with k_*t*_ *values*

For proteins whose *k*_*t*_ values passed coverage and goodness-of-fit criteria described above, protein sequence features were evaluated as follows. Hydrophobicity was quantified by grand average of hydropathy (GRAVY) score^72^. Molar abundance of amino acid classes, isoelectric point, and molecular weight were extracted using Pepstats^73^ (https://www.ebi.ac.uk/Tools/seqstats/emboss_pepstats/). The correlation of each of these parameters to *k*_*t*_ was evaluated by calculating the Spearman correlation coefficient. Intrinsically disordered regions (IDRs) were defined by identifying stretches of at least 40 amino acids having IUPRED2^74^ disorder scores > 0.5; IDRs of at least 40 amino acids in length have been previously shown to correlate with shorter protein lifetimes in cultured cells and in yeast^11,12^. A validated list of mouse IDPs was sourced from the DisPROT database^37^.

#### Determination and analysis of cell-cycle corrected k_*deg*_ *values*

Cell-cycle-corrected protein *k*_*deg*_ values were determined by subtracting the cell doubling time *(k*_*div*_) for each tissue from the apparent protein turnover rate *(k*_*t*_) determined in that tissue. In the intestine, a significant proportion of the proteome had *Kdeg* rates very similar to *k*_*div*_. Gene ontology analyses of this subset of ∼400 proteins was performed to identify biological processes (GO:BP) and cellular components (GO:CC) that were over-represented, using Gprofiler^70^. Redundant GO terms were filtered using ReVIGO^71^, and a bubble plot of significance of enrichment vs. similarity (semantic space) was generated using Prism (GraphPad), where bubble sizes correspond to the number of proteins mapped to a term. To analyze trends in turnover for intrinsically disordered proteins (IDPs), a list of curated and experimentally validated IDPs from the DisProt database^37^ was cross-referenced to *k*_*deg*_ values.

#### Cross-tissue dispersion of k_deg_ values

Cross-tissue dispersion *(D)* was calculated on a protein-by-protein basis for all proteins detected in the liver, heart, and fat tissues. *D = variance / mean*, where variance (V) is the average of the squared differences of each *k*_*deg*_ value from the mean *k*_*deg*_ value. Because *D* is normalized to the mean *k*_*deg*_ value, it is independent of the magnitude of *k*_*deg*_ (see Supplementary Figure 11). *D* values are reported in Supplementary Table 7. Analysis of *D* by organelle was performed using annotations from MitoCarta^75^ for mitochondrial proteins, from a recent proximity labeling study for lipid droplets^76^, and manually curated annotations from UniProt for all other organelles. Only UniProt annotations that listed a specific organelle as the first affiliation were retained to limit multi-localizing proteins.

#### Intra-complex dispersion of k_deg_ values for multiprotein complexes

A mouse proteome multiprotein complex subunit annotation set from ComplexPortal^77^ was used to search for multiprotein complexes with at least 5 subunits detected in the liver, heart, intestine, or fat. Intra-complex dispersion for these complexes was determined by calculating the dispersion *(d = variance / mean)* of *k*_*deg*_ values for all subunits detected in a tissue.

#### Determination of relative protein abundance within tissues

To evaluate relative protein abundance within tissues, technical replicate unlabeled wild type (WT) samples from each multiplexed TMT run were first channel normalized, then the geometric mean was calculated to determine mean normalized intensities for each biological replicate. Protein abundance was then length-normalized by dividing each protein’s normalized intensity by the number of amino acids. Finally, samples were normalized for comparison across biological replicates by normalizing each channel to the maximum value detected. The geometric mean abundance was calculated by determining the geometric mean of the length- and channel-normalized protein abundance. These relative abundance values were used to explore the relationship between protein abundance and protein half-life (Fig. 4; Supplementary Fig. 4).

## Supporting information

Merged Supplementary Figures

## Data Availability

LC-MS/MS data have been deposited in the ProteomeXchange Consortium via the PRIDE partner repository, accessible at www.ebi.ac.uk/pride. The accession codes will be included in the final manuscript.

## Author Contributions

A.B. and S.G. conceptualized the TRAIL approach. J.Hasper performed in vivo labeling experiments, tissue isolations, and protein and DNA extractions. J. Hryhorenko and K. W. prepared and ran samples for LC-MS/MS. A.B., J.Hasper, K.W., and S.G. analyzed data. A.B. and S.G. prepared the figures. A.B., J. Hasper, and S.G. wrote the manuscript.

## Acknowledgements

We would like to acknowledge the Progeria Research Foundation (A.B.), the Chan Zuckerberg Biohub (A.B.), and the National Institutes of Health (R35 GM119502 and S10 OD025242, S.G.) for funding support. We thank Biao Wang, Esther Paolo Mirasol, Balyn Zaro, Regan Volk, Carlos Lizama Valenzuela, Yana Blokhina, Galina Schmunk, and Gabrielle Servito for assistance with tissue dissections.

## References

1. Taylor, R. C. & Dillin, A. Aging as an Event of Proteostasis Collapse. Csh Perspect Biol 3, a004440 (2011).

2. Koyuncu, S. et al. Rewiring of the ubiquitinated proteome determines ageing in C. elegans. Nature 596, 285–290 (2021).

3. Price, J. C., Guan, S., Burlingame, A., Prusiner, S. B. & Ghaemmaghami, S. Analysis of proteome dynamics in the mouse brain. Proc National Acad Sci 107, 14508–14513 (2010).

4. Savas, J. N., Toyama, B. H., Xu, T., III, J. R. Y. & Hetzer, M. W. Extremely Long-Lived Nuclear Pore Proteins in the Rat Brain. Science 335, 942–942 (2012).

5. Toyama, B. H. et al. Identification of Long-Lived Proteins Reveals Exceptional Stability of Essential Cellular Structures. Cell 154, 971–982 (2013).

6. Fornasiero, E. F. et al. Precisely measured protein lifetimes in the mouse brain reveal differences across tissues and subcellular fractions. Nature Communications 9, 71–17 (2018).

7. Mathieson, T. et al. Systematic analysis of protein turnover in primary cells. Nature Communications 1–10 (2018).

8. Taylor, A. & Davies, K. J. A. Protein oxidation and loss of protease activity may lead to cataract formation in the aged lens. Free Radical Bio Med 3, 371–377 (1987).

9. D’Angelo, M. A., Raices, M., Panowski, S. H. & Hetzer, M. W. Age-Dependent Deterioration of Nuclear Pore Complexes Causes a Loss of Nuclear Integrity in Postmitotic Cells. Cell 136, 284–295 (2009).

10. Marrero, M. C. & Barrio-Hernandez, I. Toward Understanding the Biochemical Determinants of Protein Degradation Rates. Acs Omega 6, 5091–5100 (2021).

11. Lee, R. van der et al. Intrinsically Disordered Segments Affect Protein Half-Life in the Cell and during Evolution. Cell Reports 8, 1832–1844 (2014).

12. Fishbain, S. et al. Sequence composition of disordered regions fine-tunes protein half-life. Nat Struct Mol Biol 22, 214–221 (2015).

13. Wu, C. et al. Global and Site-Specific Effect of Phosphorylation on Protein Turnover. Dev Cell 56, 111-124.e6 (2021).

14. Zecha, J. et al. Linking post-translational modifications and protein turnover by site-resolved protein turnover profiling. Nat Commun 13, 165 (2022).

15. Mallik, S. & Kundu, S. Topology and Oligomerization of Mono- and Oligomeric Proteins Regulate Their Half-Lives in the Cell. Structure 26, 869-878.e3 (2018).

16. Dörrbaum, A. R., Kochen, L., Langer, J. D. & Schuman, E. M. Local and global influences on protein turnover in neurons and glia. eLife 7, 489 (2018).

17. Swovick, K. et al. Cross-species Comparison of Proteome Turnover Kinetics*. Journal Title Molecular and Cellular Proteomics 17, 580–591 (2020).

18. Matsuda, M. et al. Species-specific segmentation clock periods are due to differential biochemical reaction speeds. Science 369, 1450–1455 (2020).

19. Rolfs, Z. et al. An atlas of protein turnover rates in mouse tissues. Nat Commun 12, 6778 (2021).

20. McClatchy, D. B., Dong, M.-Q., Wu, C. C., Venable, J. D. & Yates, J. R. 15N Metabolic Labeling of Mammalian Tissue with Slow Protein Turnover. J Proteome Res 6, 2005–2010 (2007).

21. Reome, J. B. et al. The Effects of Prolonged Administration of 5-Bromodeoxyuridine on Cells of the Immune System. J Immunol 165, 4226–4230 (2000).

22. Asher, E., Payne, C. M. & Bernstein, C. Evaluation of Cell Death in EB V-Transformed Lymphocytes Using Agarose Gel Electrophoresis, Light Microscopy and Electron Microscopy: II. Induction of Non-Classic Apoptosis (“Para-Apoptosis”) by Tritiated Thymidine. Leukemia Lymphoma 19, 107–119 (2009).

23. Neese, R. A. et al. Measurement in vivo of proliferation rates of slow turnover cells by 2H2O labeling of the deoxyribose moiety of DNA. Proceedings of the National Academy of Sciences 99, 15345–15350 (2002).

24. Thompson, A. C. S. et al. Reduced in vivo hepatic proteome replacement rates but not cell proliferation rates predict maximum lifespan extension in mice. Aging Cell 15, 118–127 (2016).

25. Drake, J. C. et al. Assessment of Mitochondrial Biogenesis and mTORC1 Signaling During Chronic Rapamycin Feeding in Male and Female Mice. Journals Gerontology Ser 68, 1493–1501 (2013).

26. Miller, B. F. et al. CORP: The use of deuterated water for the measurement of protein synthesis. J Appl Physiol 128, 1163–1176 (2020).

27. Welle, K. A. et al. Time-resolved Analysis of Proteome Dynamics by Tandem Mass Tags and Stable Isotope Labeling in Cell Culture (TMT-SILAC) Hyperplexing*. Mol Cell Proteomics 15, 3551–3563 (2016).

28. Guan, S., Price, J. C., Ghaemmaghami, S., Prusiner, S. B. & Burlingame, A. L. Compartment Modeling for Mammalian Protein Turnover Studies by Stable Isotope Metabolic Labeling. Anal Chem 84, 4014–4021 (2012).

29. Guan, S., Price, J. C., Prusiner, S. B., Ghaemmaghami, S. & Burlingame, A. L. A Data Processing Pipeline for Mammalian Proteome Dynamics Studies Using Stable Isotope Metabolic Labeling*. Mol Cell Proteomics 10, M111.010728 (2011).

30. Ross, A. B., Langer, J. D. & Jovanovic, M. Proteome Turnover in the Spotlight: Approaches, Applications, and Perspectives. Journal Title Molecular and Cellular Proteomics 20, 100016 (2021).

31. Sender, R. & Milo, R. The distribution of cellular turnover in the human body. Nature Medicine 1–15 (2021).

32. Lau, E. et al. A large dataset of protein dynamics in the mammalian heart proteome. Scientific Data 3, 160015–15 (2016).

33. Krishna, S. et al. Identification of long-lived proteins in the mitochondria reveals increased stability of the electron transport chain. Dev Cell (2021) doi:10.1016/j.devcel.2021.10.008.

34. Bomba-Warczak, E., Edassery, S. L., Hark, T. J. & Savas, J. N. Long-lived mitochondrial cristae proteins in mouse heart and brain. J Cell Biol 220, e202005193 (2021).

35. Martin-Perez, M. & Villén, J. Determinants and Regulation of Protein Turnover in Yeast. Cell Syst 5, 283-294.e5 (2017).

36. Marrero, M. C., Dijk, A. D. J. van & Ridder, D. de. Sequence-based analysis of protein degradation rates. Proteins Struct Funct Bioinform 85, 1593–1601 (2017).

37. Quaglia, F. et al. DisProt in 2022: improved quality and accessibility of protein intrinsic disorder annotation. Nucleic Acids Res 50, D480–D487 (2021).

38. Drigo, R. A. e et al. Age Mosaicism across Multiple Scales in Adult Tissues. Cell Metab 30, 343-351.e3 (2019).

39. Liao, Q., Chiu, N. H. L., Shen, C., Chen, Y. & Vouros, P. Investigation of Enzymatic Behavior of Benzonase/Alkaline Phosphatase in the Digestion of Oligonucleotides and DNA by ESI-LC/MS. Anal Chem 79, 1907–1917 (2007).

40. Reichard, P. Interactions Between Deoxyribonucleotide And DNA Synthesis. Annu Rev Biochem 57, 349–374 (1988).

41. Macallan, D. C. et al. Measurement of cell proliferation by labeling of DNA with stable isotope-labeled glucose: studies in vitro, in animals, and in humans. P Natl Acad Sci Usa 95, 708–13 (1998).

42. Quinlivan, E. P. & III, J. F. G. DNA digestion to deoxyribonucleoside: A simplified one-step procedure. Analytical Biochemistry 373, 383–385 (2008).

43. Darwich, A. S., Aslam, U., Ashcroft, D. M. & Rostami-Hodjegan, A. Meta-Analysis of the Turnover of Intestinal Epithelia in Preclinical Animal Species and Humans. Drug Metab Dispos 42, 2016–2022 (2014).

44. MacDONALD, R. A. Lifespan of Liver Cells: Autoradiographic Study Using Tritiated Thymidine in Normal, Cirrhotic, and Partially Hepatectomized Rats. Arch Intern Med 107, 335–343 (1961).

45. Malliaras, K. et al. Cardiomyocyte proliferation and progenitor cell recruitment underlie therapeutic regeneration after myocardial infarction in the adult mouse heart. Embo Mol Med 5, 191–209 (2013).

46. Rigamonti, A., Brennand, K., Lau, F. & Cowan, C. A. Rapid Cellular Turnover in Adipose Tissue. Plos One 6, e17637 (2011).

47. Wu, R. S., Tsai, S. & Bonner, W. M. Patterns of histone variant synthesis can distinguish go from G1 cells. Cell 31, 367–374 (1982).

48. Ahmad, K. & Henikoff, S. The Histone Variant H3.3 Marks Active Chromatin by Replication-Independent Nucleosome Assembly. Mol Cell 9, 1191–1200 (2002).

49. Cook, A. J. L., Gurard-Levin, Z. A., Vassias, I. & Almouzni, G. A Specific Function for the Histone Chaperone NASP to Fine-Tune a Reservoir of Soluble H3-H4 in the Histone Supply Chain. Mol Cell 44, 918–927 (2011).

50. Mizushima, N., Yamamoto, A., Matsui, M., Yoshimori, T. & Ohsumi, Y. In vivo analysis of autophagy in response to nutrient starvation using transgenic mice expressing a fluorescent autophagosome marker. Molecular Biology of the Cell 15, 1101–1111 (2004).

51. Vonk, W. I. M. et al. Differentiation Drives Widespread Rewiring of the Neural Stem Cell Chaperone Network. Mol Cell 78, 329-345.e9 (2020).

52. Jenkins, E. C. et al. Proteasome mapping reveals sexual dimorphism in tissue-specific sensitivity to protein aggregations. Embo Rep 21, e48978 (2020).

53. Jr, W. A. D. et al. Pexophagy: The Selective Autophagy of Peroxisomes. Autophagy 1, 75–83 (2005).

54. Singh, R. et al. Autophagy regulates lipid metabolism. Nature 458, 1131–1135 (2009).

55. Youle, R. J. & Narendra, D. P. Mechanisms of mitophagy. Nat Rev Mol Cell Bio 12, 9–14 (2011).

56. Monastyrska, I. & Klionsky, D. J. Autophagy in organelle homeostasis: Peroxisome turnover. Mol Aspects Med 27, 483–494 (2006).

57. Zechner, R., Madeo, F. & Kratky, D. Cytosolic lipolysis and lipophagy: two sides of the same coin. Nat Rev Mol Cell Bio 18, 671–684 (2017).

58. Lafontan, M. & Langin, D. Lipolysis and lipid mobilization in human adipose tissue. Prog Lipid Res 48, 275–297 (2009).

59. Pascual, F. & Coleman, R. A. Fuel availability and fate in cardiac metabolism: A tale of two substrates. Biochimica Et Biophysica Acta Bba - Mol Cell Biology Lipids 1861, 1425–1433 (2016).

60. McWilliams, T. G. et al. mito-QC illuminates mitophagy and mitochondrial architecture in vivo. J Cell Biol 214, 333–345 (2016).

61. Winter, D. & Becker, T. Surveying the mitochondrial proteome. Nat Cell Biol 23, 1216–1217 (2021).

62. Altshuler-Keylin, S. et al. Beige Adipocyte Maintenance Is Regulated by Autophagy-Induced Mitochondrial Clearance. Cell Metab 24, 402–419 (2016).

63. McShane, E. et al. Kinetic Analysis of Protein Stability Reveals Age-Dependent Degradation. Cell 167, 803-815.e21 (2016).

64. Matera, A. G. & Wang, Z. A day in the life of the spliceosome. Nat Rev Mol Cell Bio 15, 108–121 (2014).

65. Bussiere, C., Hashem, Y., Arora, S., Frank, J. & Johnson, A. W. Integrity of the P-site is probed during maturation of the 60S ribosomal subunit. J Cell Biol 197, 747–759 (2012).

66. Shi, Z. et al. Heterogeneous Ribosomes Preferentially Translate Distinct Subpools of mRNAs Genome-wide. Mol Cell 67, 71-83.e7 (2017).

67. Neff, N. F. et al. Single-cell transcriptomics of 20 mouse organs creates a Tabula Muris. Nature 1–25 (2018).

68. McAlister, G. C. et al. MultiNotch MS3 Enables Accurate, Sensitive, and Multiplexed Detection of Differential Expression across Cancer Cell Line Proteomes. Anal Chem 86, 7150–7158 (2014).

69. Su, D. et al. Quantitative analysis of ribonucleoside modifications in tRNA by HPLC-coupled mass spectrometry. Nat Protoc 9, 828–841 (2014).

70. Raudvere, U. et al. g:Profiler: a web server for functional enrichment analysis and conversions of gene lists (2019 update). Nucleic Acids Res 47, W191–W198 (2019).

71. Supek, F., Bošnjak, M., Škunca, N. & Šmuc, T. REVIGO Summarizes and Visualizes Long Lists of Gene Ontology Terms. Plos One 6, e21800 (2011).

72. Kyte, J. & Doolittle, R. F. A simple method for displaying the hydropathic character of a protein. J Mol Biol 157, 105–132 (1982).

73. Madeira, F. et al. Search and sequence analysis tools services from EMBL-EBI in 2022. Nucleic Acids Res 50, W276–W279 (2022).

74. Mészáros, B., Erdős, G. & Dosztányi, Z. IUPred2A: context-dependent prediction of protein disorder as a function of redox state and protein binding. Nucleic Acids Res 46, W329–W337 (2018).

75. Rath, S. et al. MitoCarta3.0: an updated mitochondrial proteome now with sub-organelle localization and pathway annotations. Nucleic Acids Res 49, gkaa1011.(2020).

76. Bersuker, K. et al. A Proximity Labeling Strategy Provides Insights into the Composition and Dynamics of Lipid Droplet Proteomes. Dev Cell 44, 97-112.e7 (2018).

77. Meldal, B. H. M. et al. Complex Portal 2018: extended content and enhanced visualization tools for macromolecular complexes. Nucleic Acids Res 47, gky1001.(2018).

