## Supplementary material for "Turnover and replication analysis by isotope labeling (TRAIL) reveals the influence of tissue context on protein and organelle lifetimes": Merged Supplementary Figures

Supplementary Figure 1

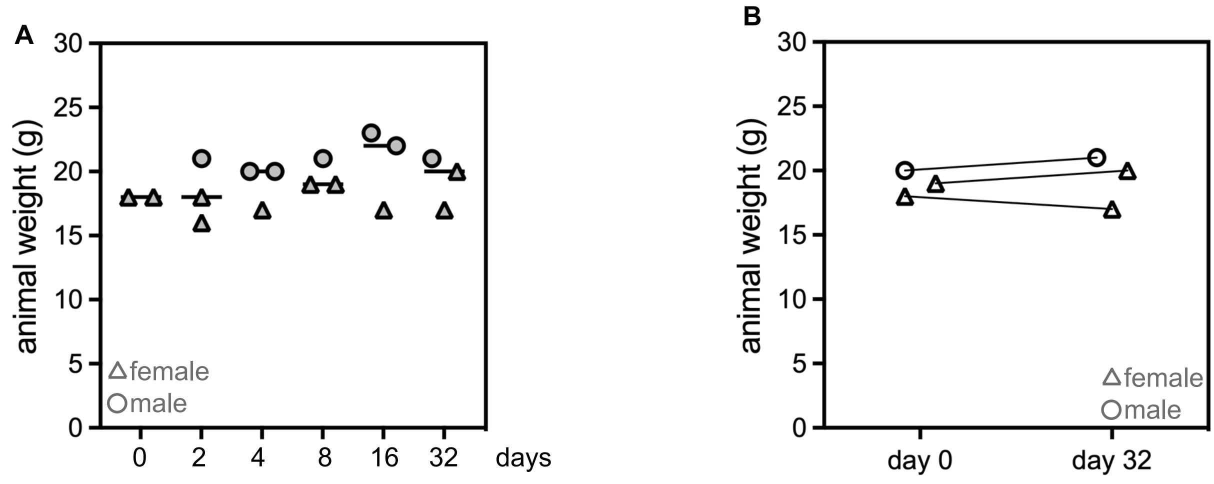


Supplementary Figure 1. (A) Weights of animals on date of sacrifice for TRAIL timecourse. (B) Weight of 3 animals over 32 days of labeling.

A B

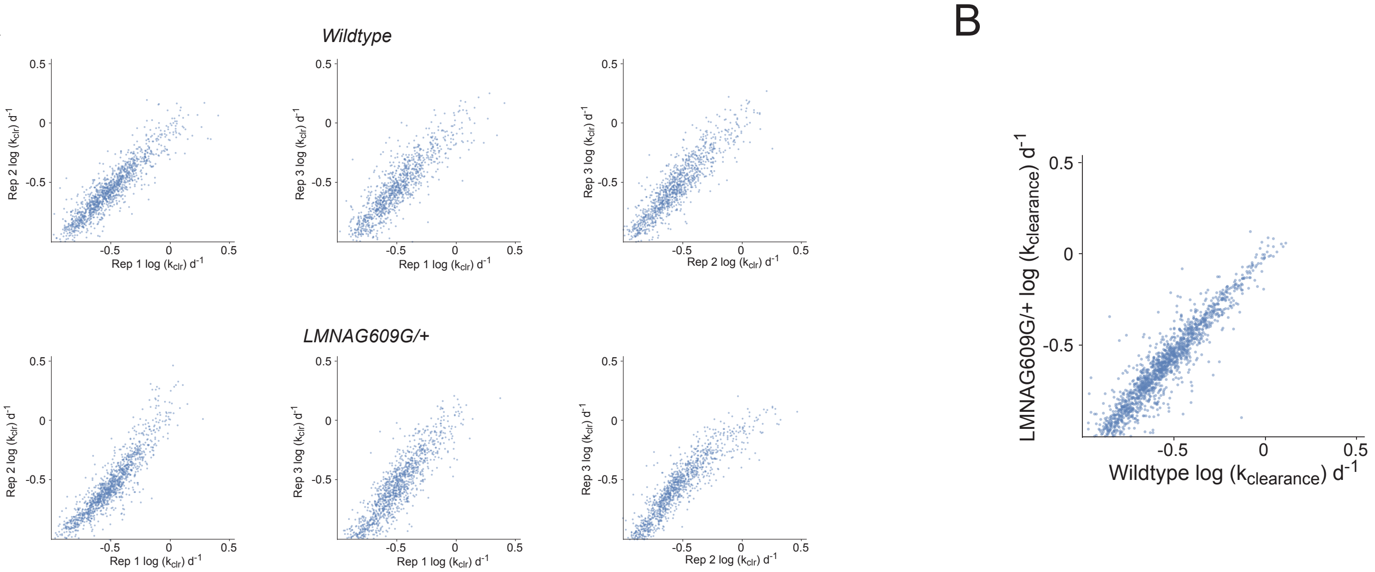


C

| | proteins detected | proteins quantified | median PSMs quantified per protein | median $k_{clr}$ (d-1) | median $k_{clr}$ std. error(d-1) |
|---|---|---|---|---|---|
| liver | 3,390 | 1,577 | 21.5 | 0.282 | 0.016 |
| heart | 2,652 | 1,187 | 24.6 | 0.106 | 0.007 |
| fat | 2,875 | 1,104 | 23.0 | 0.115 | 0.010 |
| intestine | 4,329 | 2,019 | 22.5 | 0.395 | 0.026 |

D

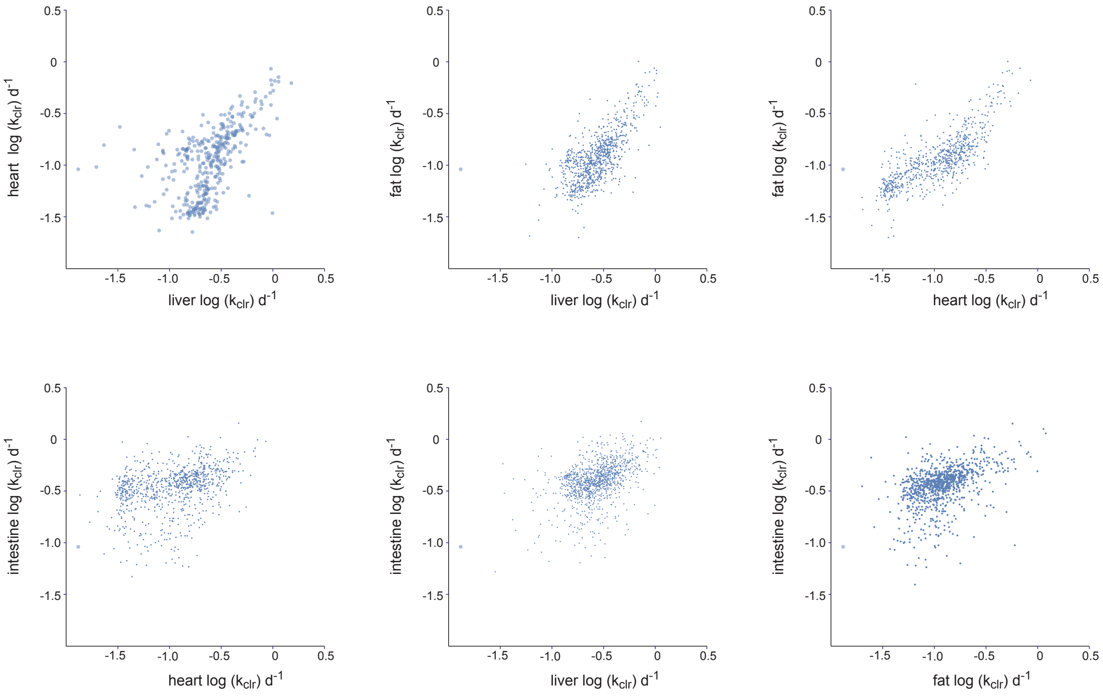

Supplementary Figure 2. (A) Representative scatterplots showing high concordance of $k_t$ values determined across 3 replicate experiments each in wild type (top) and progeroid (*LMNA*$^{G609G/+}$, bottom) liver. (B) Scatterplot showing correlation in $k_t$ values between wild type and progeroid liver. (C) Overview of proteins detected and quantified in each tissue. (D) Pairwise correlation of $k_t$ values for proteins detected in liver, heart, fat, and intestine.

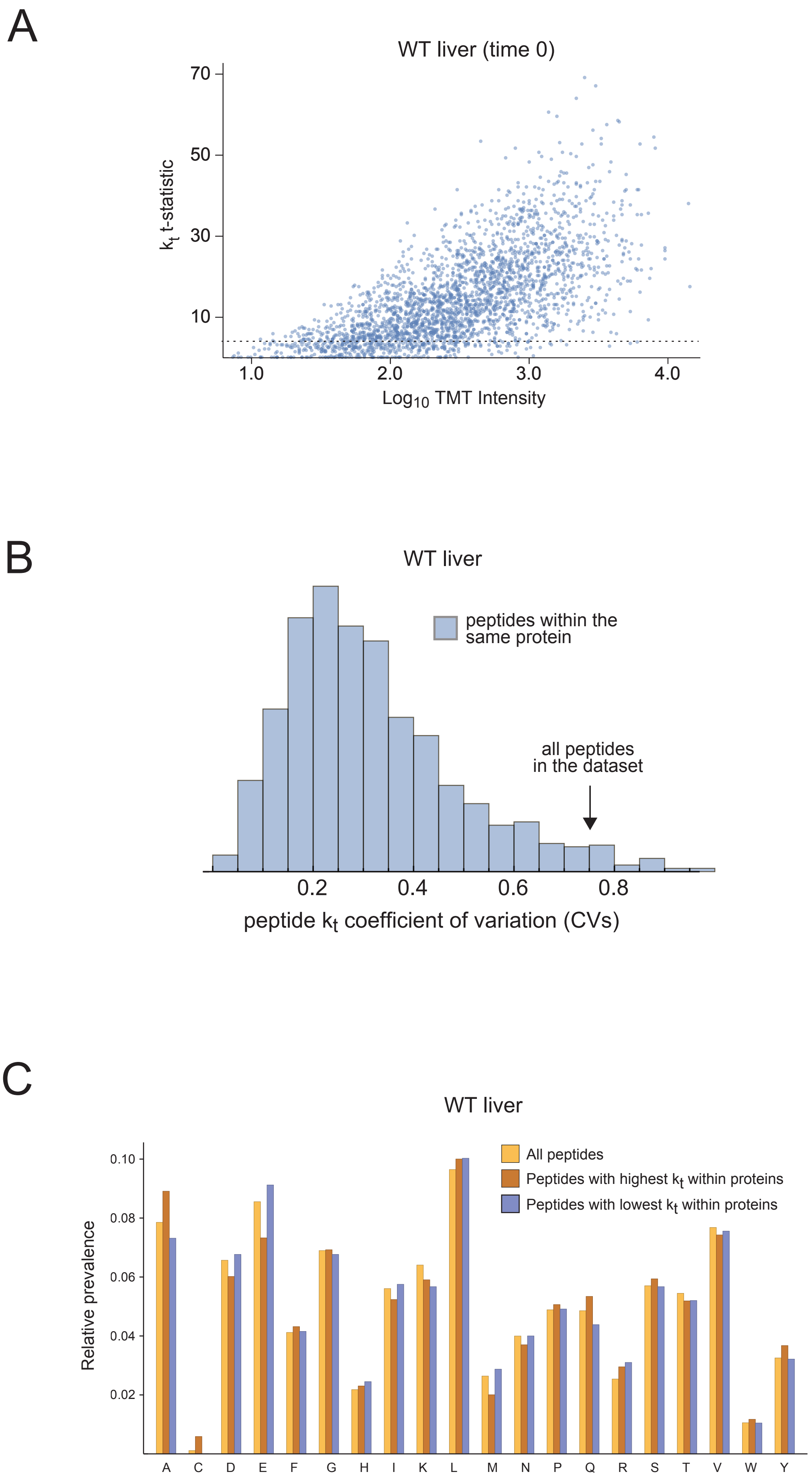
A
WT liver (time 0)
$k_t$ t-statistic
70
50
30
10
1.0
2.0
3.0
4.0
Log$_{10}$ TMT Intensity
B
WT liver
peptides within the same protein
all peptides in the dataset
0.2
0.4
0.6
0.8
peptide $k_t$ coefficient of variation (CVs)
C
WT liver
Relative prevalence
0.10
0.08
0.06
0.04
0.02
All peptides
Peptides with highest $k_t$ within proteins
Peptides with lowest $k_t$ within proteins
A
C
D
E
F
G
H
I
K
L
M
N
P
Q
R
S
T
V
W
Y

Supplementary Figure 3. (A) Higher intensity reporter ions lead to rate measurements with lower relative errors. As an example, the plot shows that the intensities of TMT reporter ions for WT liver (t=0) measurements are positively correlated with the t-statistics of $k_t$ measurements. A similar trend is observed for all of our rate measurements. Thus, as expected, higher intensity ions result in more precise $k_t$ measurements However, as described in the text, we use strict criteria (>3 PSMs, 2 or more replicate measurements and t-stat>3 as shown by the dotted line in the plot) to filter our measurements prior to downstream analyses. (B) The variabilities of $k_t$ measurements for peptides mapped to the same protein are significantly lower than the peptides in the dataset at large. This can be shown by the fact that the distribution of coefficient of variations (CVs) of peptides within the same proteins is significantly lower than all peptides in our dataset. The figure demonstrates this for WT liver data and the same trend holds for all tissues. (C) Amino acid composition of peptides does not significantly influence the magnitude of $k_t$ measurements. To demonstrate this, we analyzed the prevalence of amino acids within the fastest and slowest labeling peptides within the proteins in our dataset. As can be seen in the plot for the WT liver data, there is no significant enrichment for specific amino acids within fast or slow labeling peptides. The same trend holds for other tissues.

Supplementary Figure 4.

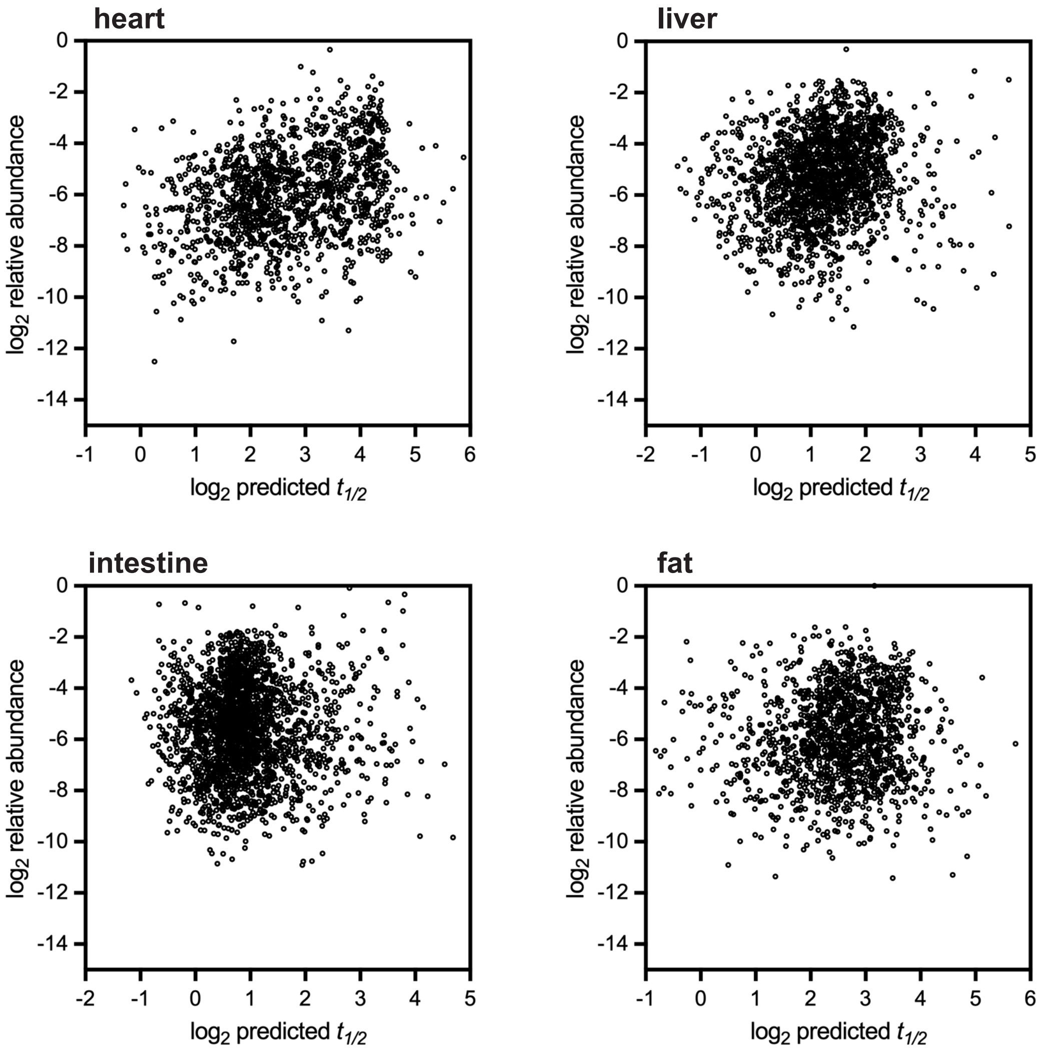


Supplementary Figure 4. Analysis of correlation between relative protein abundance and protein half-life in wild-type heart, liver, intestine and fat.

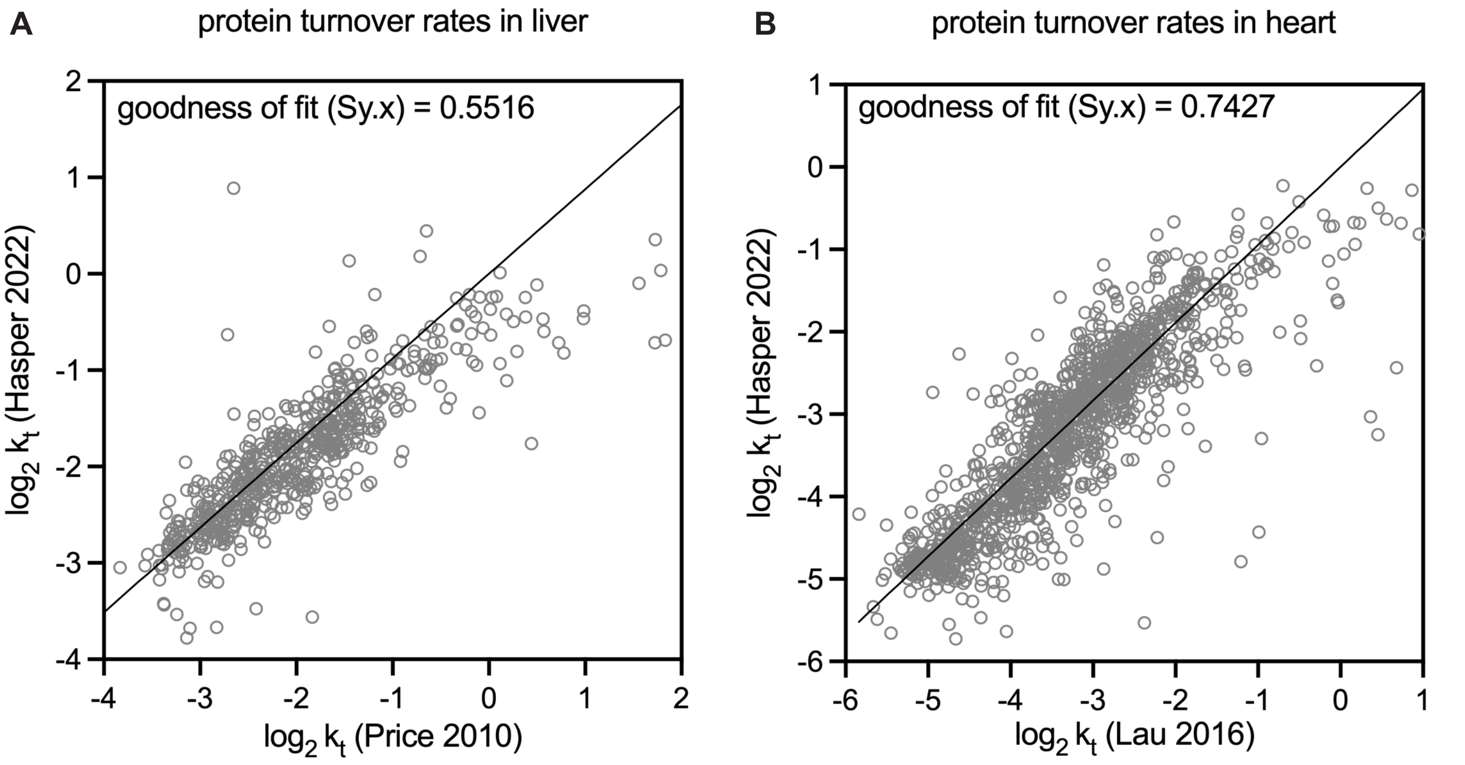


Supplementary Figure 5. Comparison of protein turnover rates determined in this study to previously published protein turnover data determined by $^{15}N$ labeling of the mouse liver (A, Price et al., *PNAS* 2010) and by $D_2O$ labeling of the mouse heart (B, Lau et al., *Sci Data* 2016).

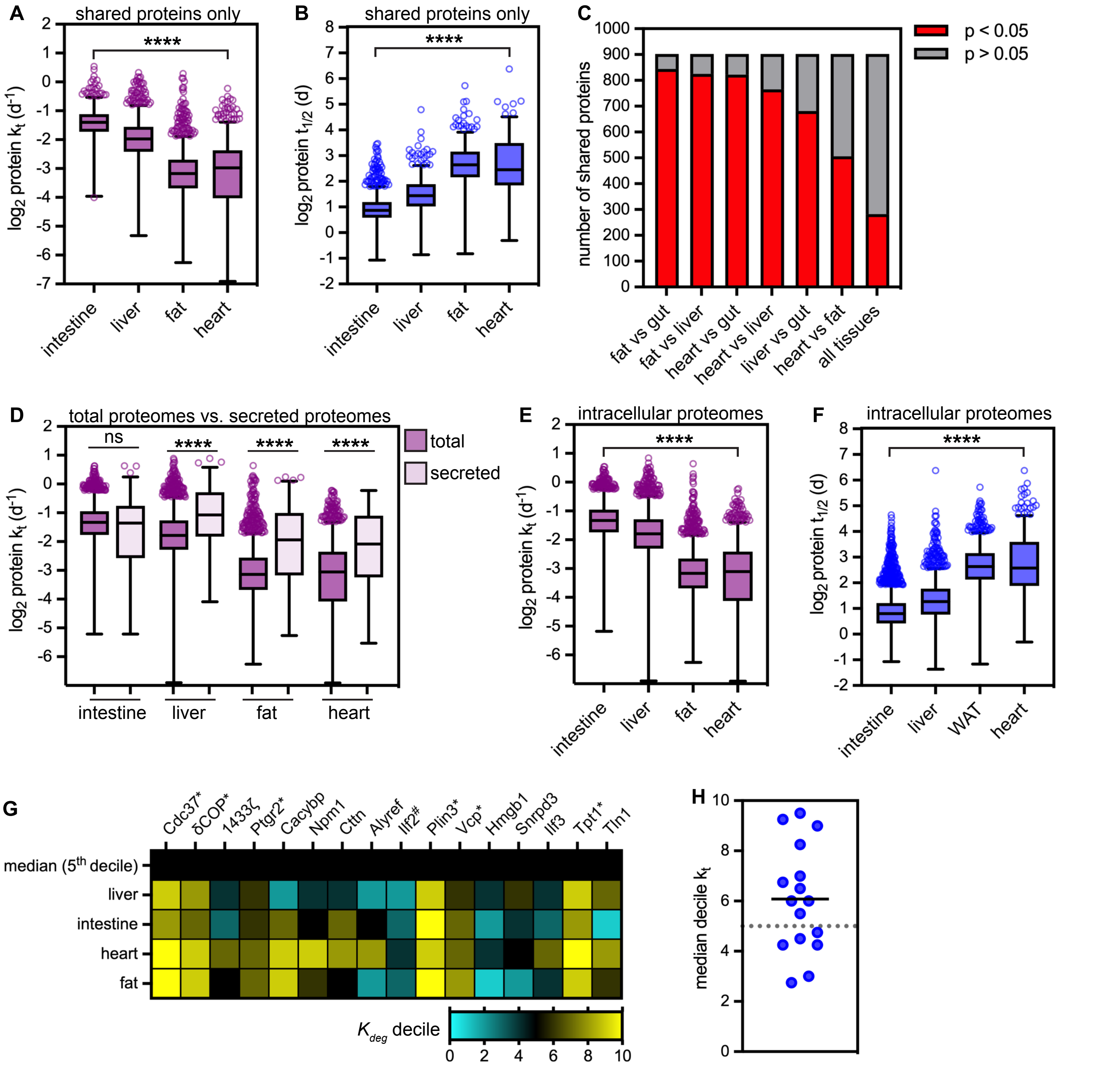


**Supplementary Figure 6**. (A) Degradation rates ($k_t$) and (B) predicted half-lives ($t_{1/2}$) determined by SILAM-TMT of 902 proteins detected across intestine, liver, fat, and heart. **** indicates that all medians are significantly different ($p < 0.0001$, Kruskal-Wallis test). Median $t_{1/2}$ for the shared proteomes of these four tissues are 1.8 days (intestine), 2.7 days (liver), 6.3 days (fat), and 5.5 days (heart). (C) Proportion of proteins whose values of $k_t$ are significantly different across the indicated pairwise comparisons. Significance determined by t-test. (D) Comparison of turnover rates of secreted proteins (149 in intestine; 103 in liver; 99 in fat; and 96 in heart) versus total tissue proteomes (data reproduced from Figure 3A). **** indicates that distributions are significantly different in the heart, liver, and fat ($p < 0.0001$, Mann-Whitney test), but not in the intestine. (E) $k_t$ and (F) predicted $t_{1/2}$ for intracellular proteomes. **** indicates that all medians are significantly different ($p < 0.0001$, Kruskal-Wallis test). Median half-lives for the intracellular proteomes of these four tissues are 1.7 days (intestine, n = 2570), 2.4 days (liver, n = 1996), 6.2 days (fat, n = 1511), and 6 days (heart, n = 1539). (G) Heatmap of $k_t$ deciles for 16 intrinsically disordered proteins (IDPs). *indicates proteins whose $k_t$ rates are significantly faster than the proteome median across all tissues; #indicates proteins whose $k_t$ rates are significantly slower than the proteome median across all tissues. IDP annotations from the DisProt database. (H) Median decile $k_t$ value for 16 IDPs across 4 tissues is not significantly different from the $k_t$ median (5th decile).

Supplementary Figure 7.

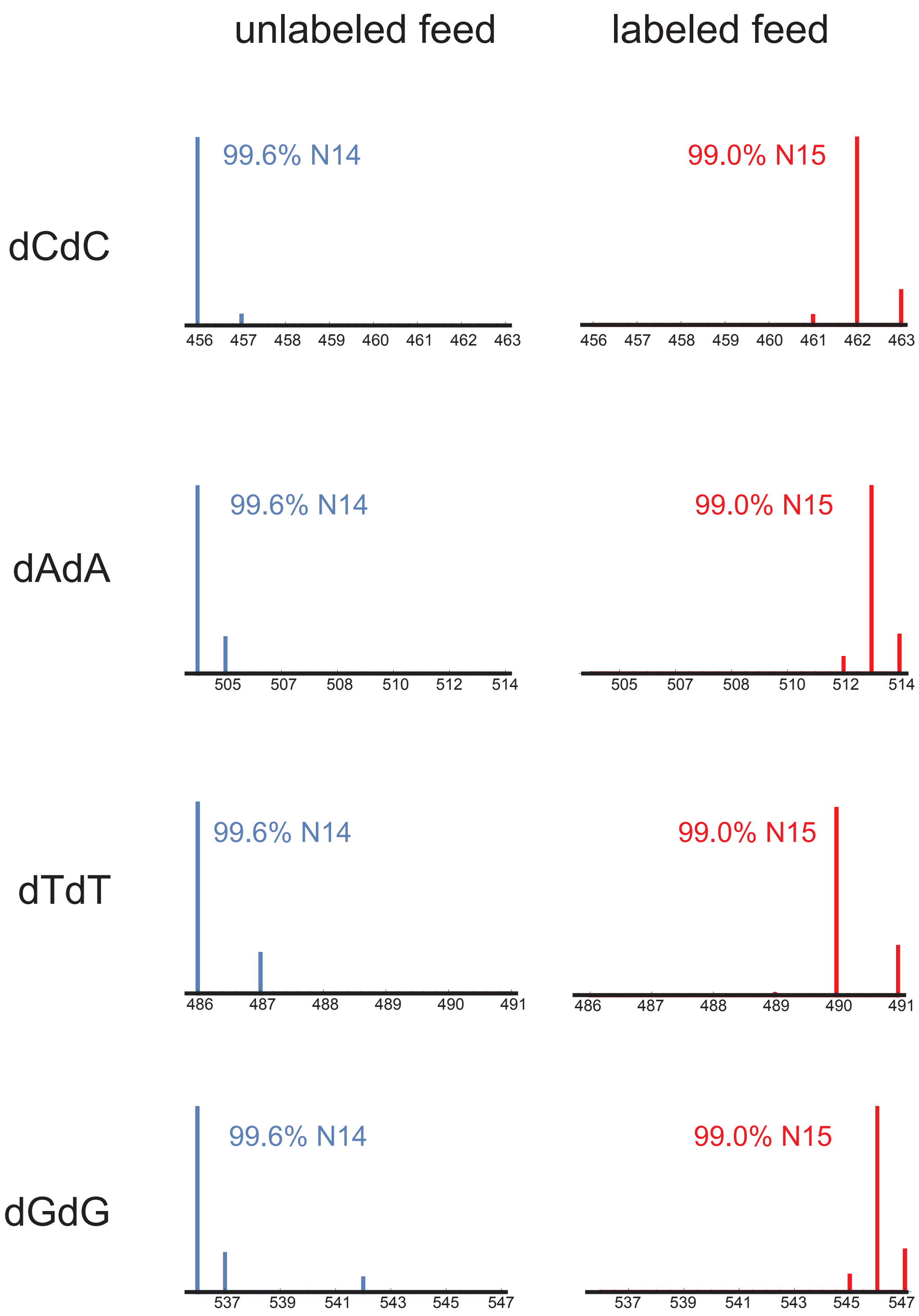


**Supplementary Figure 7.** Analysis of $^{15}N$ vs. $^{14}N$ isotope abundance in dinucleotides isolated from unlabeled chow versus $^{15}N$-labeled SILAM chow.

Supplementary Figure 8.

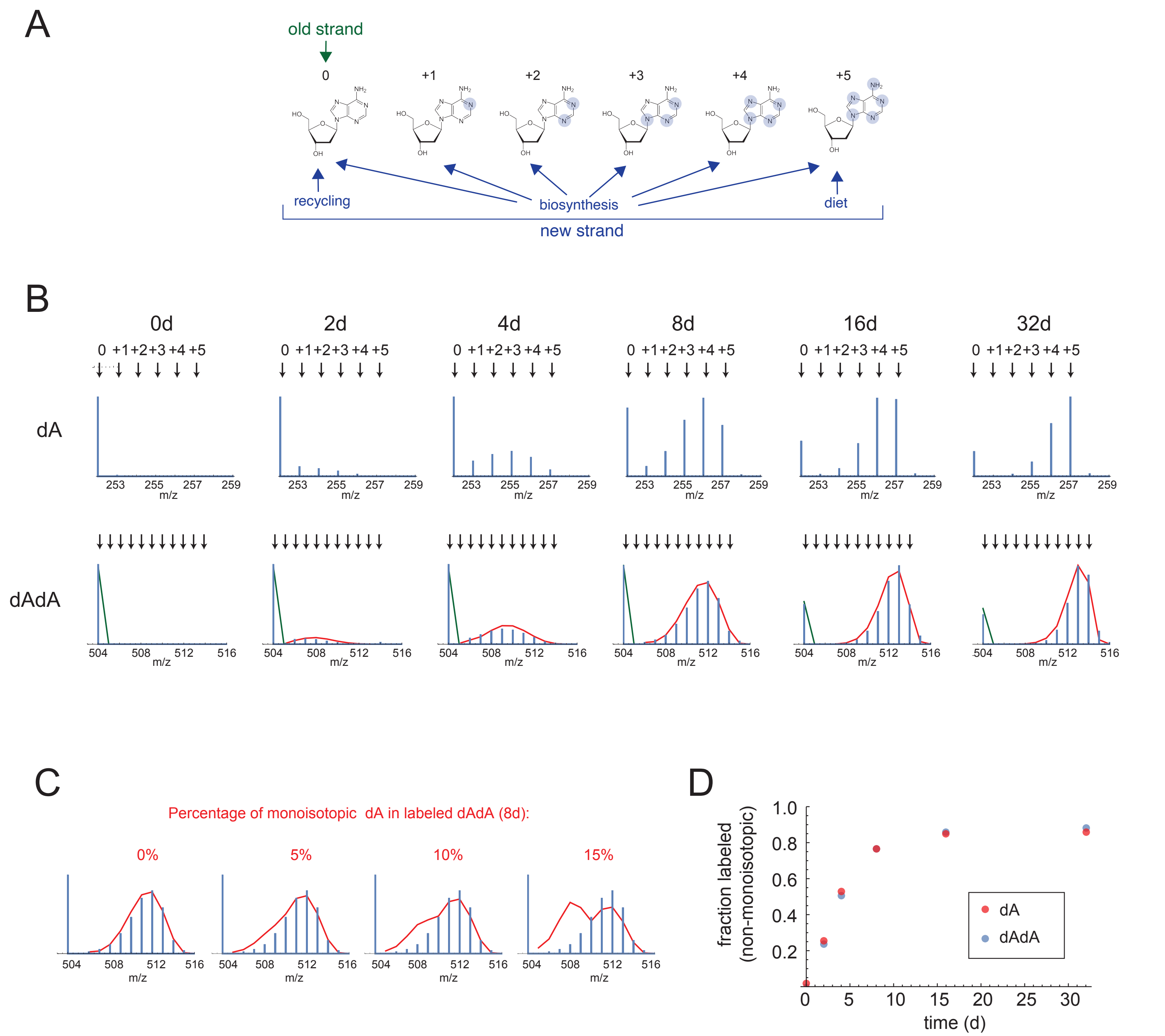


**Supplementary Figure 8**. (A) Diagram of sources of nucleotides that may be used in vivo for genomic DNA synthesis during a SILAM experiment. (B) Example labeling pattern for deoxyadenosine (dA, top) and dAdA dinucleotides (bottom) over 32 days of labeling in mouse intestine; note that the unlabeled peak is monoisotopic while the labeled peaks gradually increase in *m/z* over time. Red line indicates the distribution of labeled peaks that would be expected if the precursor pool were composed entirely of labeled species derived from either biosynthesis or diet as diagrammed in (A), with no contributioon from the unlabeled pool. (C) Fitting the observed distribution of dAdA by assuming 0-15% the source nucleotide comes from recycled unlabeled dA. Note that the best curve fit is obtained for the 0% recycling condition. (D) Close agreement of label incorporatioon observed between mononucleosides (dA) and dinucleotides (dAdA) over 32 days of labeling in wild type intestine.

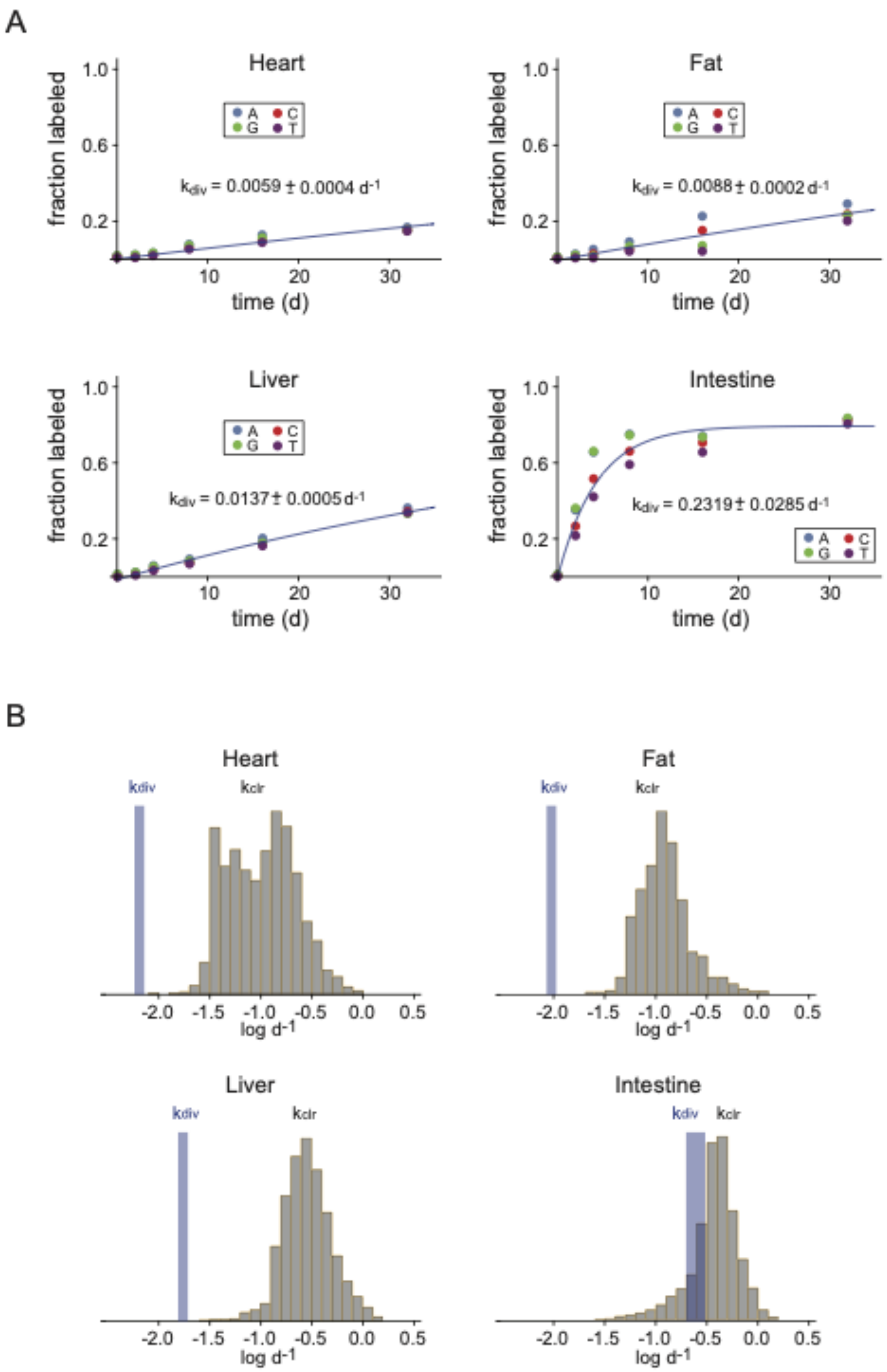


Supplementary Figure 9. (A) Curve fits for mononucleosides isolated from genomic DNA in heart, fat, liver, and intestine. (B) Comparison of proteome $k_t$ values to cell $k_{div}$ values in each tissue.

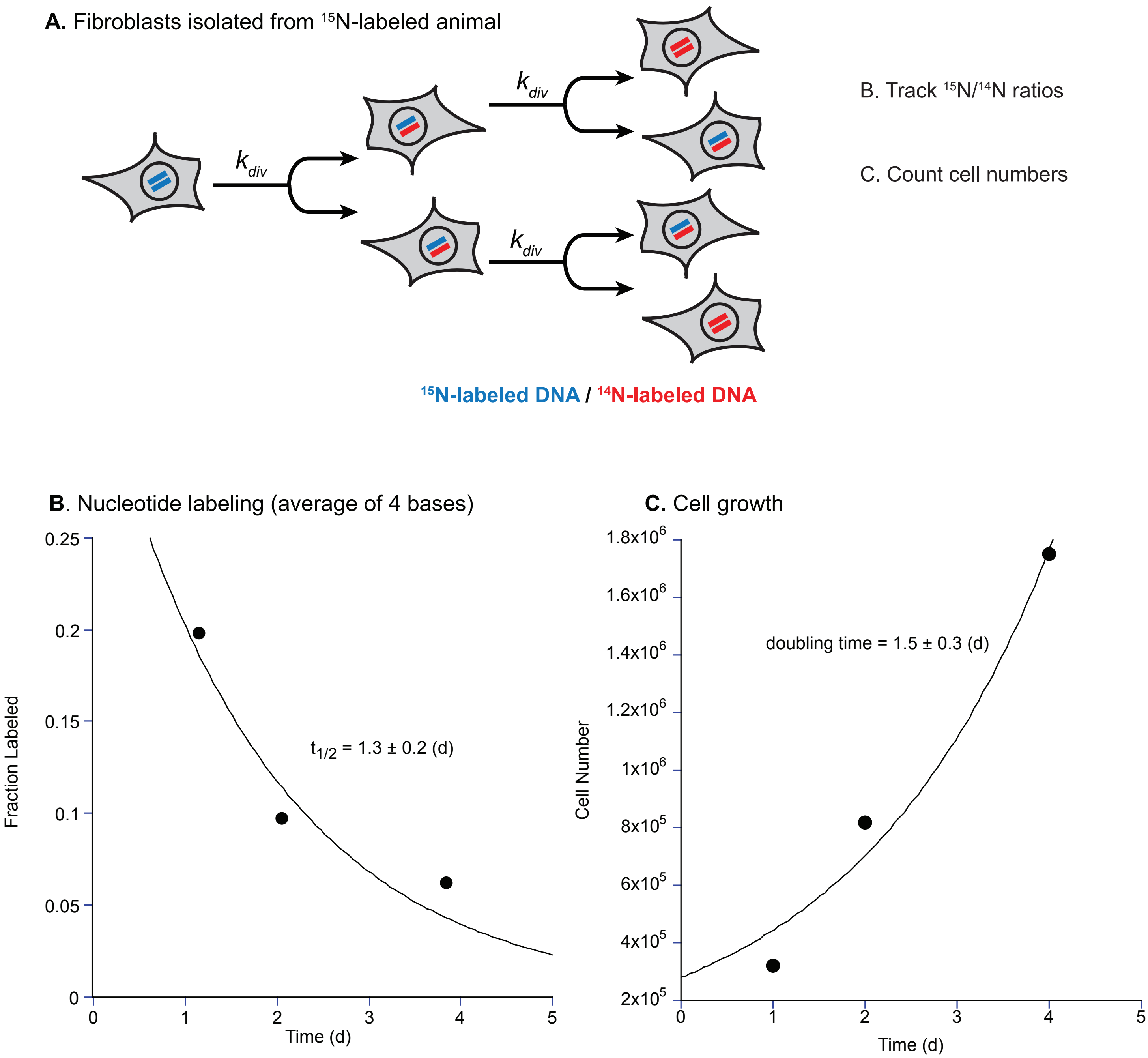


Supplementary Figure 10. (A) Schematic of cell turnover benchmarking experiment. Primary fibroblasts were isolated from the ear of a mouse labeled with 15N for 256 days. Cells were subcultured, and genomic DNA was extracted at 3 timepoints to track cell division rates by 15N/14N labeling ratios (B). At each timepoint, cell numbers were also quantified (C). Both datasets were fit to determine cell division rates and cell doubling times.

Supplementary Figure 11

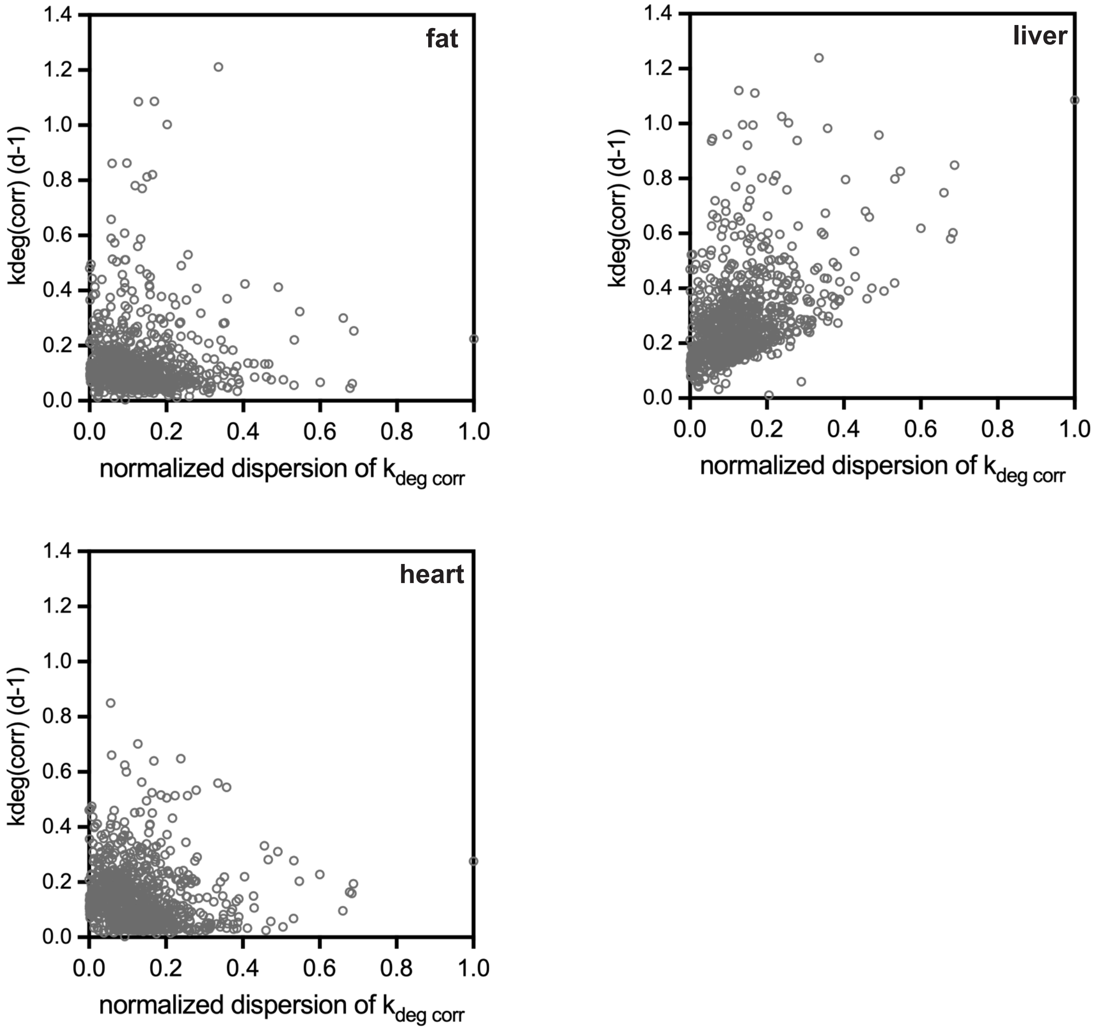


**Supplementary Figure 11**: Normalized cross-tissue dispersion of $k_{deg}$ is not correlated to the magnitude of $k_{deg}$ in each tissue.

Supplementary Figure 12.

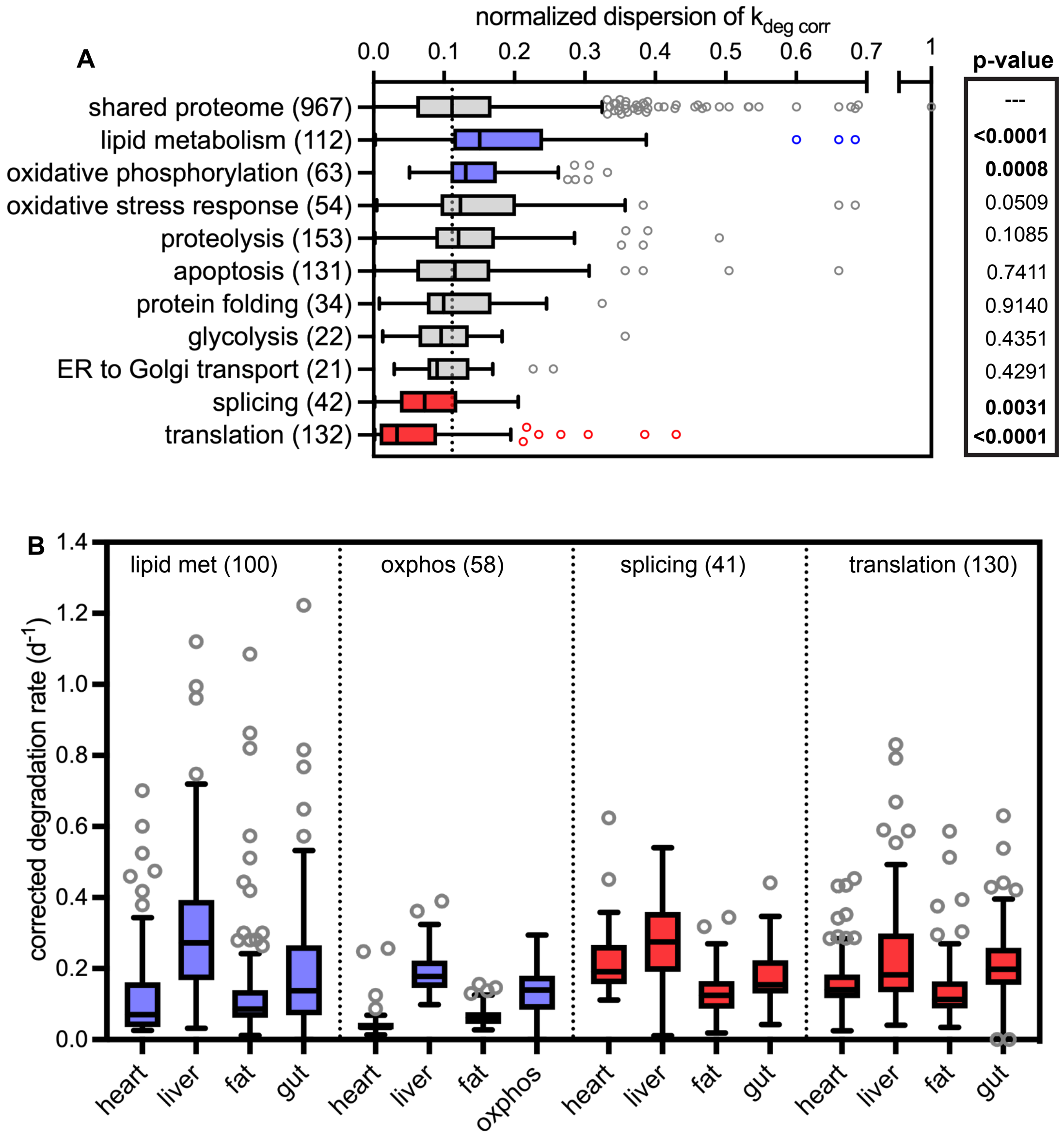


**Supplementary Figure 12**: (A) Analysis of normalized cross-tissue dispersion of $k_{deg}$ by KEGG pathway. Blue indicates metabolic pathways with components that have significantly highervariability in their degradation rates across tissues vs. the proteome median; red indicates metabolic pathways with significantly lower variability across tissues vs. the proteome median. Dotted line indicates proteome median. Significance determined by Mann-Whitney test. (B) $k_{deg}$ values of protein components of selected KEGG pathways across tissues, including proteins detected in the intestine for comparison. Blue indicates a subset has a significantly elevated median dispersion of $k_{deg}$ values across tissues; red indicates a subset has a significantly decreased dispersion of $k_{deg}$ values across tissues.

Supplementary Figure 13.

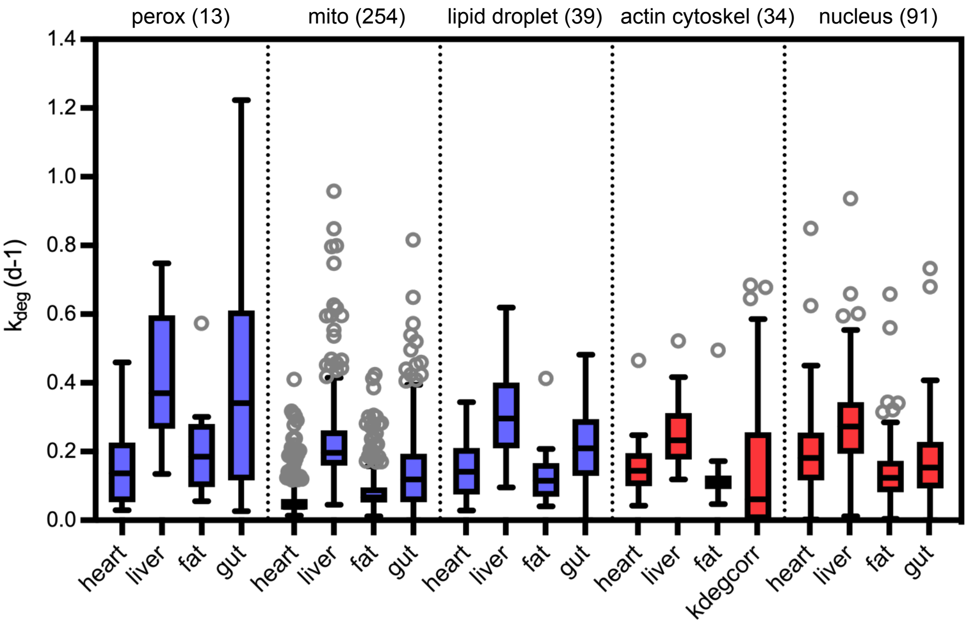


**Supplementary Figure 13**: $k_{deg}$ values of protein components of selected organelles across tissues, including proteins detected in the intestine for comparison. Blue indicates a subset has a significantly elevated median dispersion of $k_{deg}$ values across tissues; red indicates a subset has a significantly decreased dispersion of $k_{deg}$ values across tissues. See also Supplementary Table 5 for full dataset..
